# Fat taste responsiveness, but not dietary fat intake, is affected in *Adipor1* null mice

**DOI:** 10.1101/2025.03.12.642880

**Authors:** Fangjun Lin, Timothy A. Gilbertson

## Abstract

Taste is a major driving force that influences food choices and dietary intake. Adiponectin has been shown to selectively enhance cellular responses to fatty acids by mediating the activation of AMPK and translocation of CD36 in taste cells via its receptor AdipoR1. Whether *Adipor1* gene knockout affects fat taste responsiveness and dietary fat intake in animals remains unclear. In the present study, we evaluated cellular, neural, and behavioral responses to fat, as well as the dietary fat intake in global *Adipor1* knockout mice and their WT controls. Sex-specific changes in cellular and behavioral responses to fatty acid were observed in *Adipor1* knockout mice. Linoleic acid (LA)-induced calcium responsiveness appears to be reduced in taste cells from *Adipor1*-deficient males and increased in taste cells from *Adipor1*-deficient females. Brief-access taste testing revealed a loss of fat taste behavioral responsiveness in naïve *Adipor1*^-/-^ animals. Fat taste loss found in *Adipor1*^-/-^ males was restored after fat exposure and showed no significant differences in taste behavioral responses to fatty acids with WT controls in two-bottle preference and conditioned taste aversion tests. *Adipor1*^-/-^ females were found to have diminished preference for LA in two-bottle preference tests, lower intralipid/water lick ratio in a brief-access assay, and reduced avoidance for LA in conditioned taste aversion assay. Furthermore, the taste nerve responses to intralipid and the dietary fat intakes appeared to be the same between *Adipor1*^-/-^ and WT mice. In the high-fat diet feeding study, *Adipor1*^-/-^ females gained more weight, while no differences in body weight gain were found in males. Together, we show that adiponectin/AdipoR1 signaling plays crucial sex-specific roles in the modulation of fat taste and the maintenance of healthy body weight primarily by regulating energy expenditure rather than dietary fat intake in mice.

## Introduction

The gustatory system allows us to distinguish among foods to select diets with essential nutrients and avoid those containing potentially toxic substances. Although a broad range of factors influences dietary choice, taste is one primary driving force behind food consumption, determining the type and amount of food a person or animal chooses to eat (Drewnowski, 1997; Drewnowski & Monsivais, 2020). Bitter and sour tastes are generally unacceptable, especially at high concentrations, and are therefore associated with food aversion (Liman & Kinnamon, 2021; Meyerhof et al., 2005). In contrast, salty, sweet, fatty, and umami tastes are innately accepted and contribute to food palatability, meaning that the perception of these tastes in foods tends to trigger their intake (Liem & Russell, 2019; Mattes, 2021). Thus, the wide availability and high palatability of energy-dense foods in modern urban environments leading to overconsumption are major contributing factors in the growing obesity epidemic (Swinburn et al., 2011). On the other hand, what humans and animals ingest may, in turn, affect their taste sensitivities. For example, it has been shown that eating a low-fat diet makes people more sensitive to fatty acids, while excessive dietary fat significantly reduces taste sensitivity to fatty acids (Stewart & Keast, 2012). Therefore, exploring the links between taste perception, food choice, and energy intake is essential to expand our understanding of factors associated with weight control and the risk of obesity-related diseases.

Animals vary in their taste abilities due to factors such as genetics (Diószegi et al., 2019), gender (Dahir et al., 2021), age (Wang et al., 2020), nutritional (Masic & Yeomans, 2017), and hormonal status (Loper et al., 2015). There is accumulating evidence that several appetite-regulating hormones can modulate gustatory detection of fat, and this hormonal modulation of fat taste likely influences food palatability and selection, thereby altering intake of fat (Brissard et al., 2018; Calder et al., 2021; Crosson et al., 2019; Martin et al., 2012; Ullah et al., 2021). Among them, adiponectin is a key metabolic hormone predominantly released from adipose tissue that acts in the hypothalamus to regulate food intake. However, its role in appetite regulation is controversial, including stimulating food intake (Kubota et al., 2007), inhibiting food intake (Coope et al., 2008), and not affecting food intake (Qi et al., 2004). In addition to the brain, adiponectin has also been reported to target many peripheral tissues (Esmaili et al., 2020), exerting a myriad of beneficial effects in metabolic processes (Ghadge et al., 2018), and is inversely associated with insulin resistance (Li et al., 2020; Yamauchi et al., 2001), obesity (Arita et al., 1999), and hypertension (Adamczak et al., 2003). Importantly, adiponectin receptors are found to be highly expressed in mouse taste bud cells and salivary gland-specific adiponectin rescue, specifically increased behavioral taste responses to fat stimuli in adiponectin knockout mice (Crosson et al., 2019). Our more recent studies reported that adiponectin/AdipoRon selectively enhances fatty acid-induced calcium responses in immortalized human fungiform taste cells and mouse taste bud cells by increasing the surface expression of CD36 (Lin et al., 2023). This fat-enhancing effect of adiponectin/AdipoRon in taste cells is mediated by the activation of 5′ AMP-activated protein kinase (AMPK) through its receptor AdipoR1 (Lin et al., 2024). *Adipor1*-deficient male mice apparently failed to taste the intralipid solutions upon initial exposure to this tastant, while *Adipor1*-deficient female animals could detect the difference between intralipid and water but were indifferent to increasing concentrations of intralipid (Lin et al., 2024). These findings suggest that adiponectin signaling has a profound, potentially sex-dependent, function in regulating fat taste.

This study aimed to further explore the impact of *Adipor1* deficiency on the peripheral fat taste reception and determine whether it affects dietary fat intake in mice. We investigated cellular, neural, and behavioral responses to fat and other taste stimuli in global *Adipor1* gene knockout mice and their WT controls. Sex-different changes in cellular responses to fatty acid were observed in taste bud cells isolated from *Adipor1* knockout mice. Linoleic acid-induced calcium responsiveness appeared to be reduced in taste cells from *Adipor1*-deficient males and increased in taste cells from *Adipor1*-deficient females. However, we failed to observe any difference in CT nerve responses to intralipid between *Adipor1*^-/-^ and WT mice. Taste behavioral assays revealed a loss of fat taste behavioral responsiveness in naïve *Adipor1*^-/-^ animals. Fat taste loss found in *Adipor1*^-/-^ males was restored after fat exposure and showed no significant differences in taste behavioral responses to fatty acids with WT controls, whereas *Adipor1*^-/-^ females reduced gustatory behavioral responses to fatty acids in all three behavioral assays. In addition, the dietary fat intake appeared to be the same between *Adipor1*^-/-^ and WT mice, but adiponectin/AdipoR1 signaling may participate in a sex-dependent fashion in weight control in mice, especially under a high-fat diet, by regulating energy expenditure rather than dietary intake in mice.

## Results

### Calcium responses to linoleic acid in taste cells of Adipor1^-/-^ and WT mice

Our previous studies have demonstrated that AdipoRon/adiponectin could enhance fatty acid-induced calcium responses by mediating the activation of AMPK and the translocation of CD36 via *Adipor1* in taste cells (Lin et al., 2023). Here, we sought to determine whether the cellular responses to fatty acid in taste bud cells isolated from global *Adipor1* knockout mice would differ from taste bud cells from WT controls. Although no significant genotype effect was observed on LA responses between circumvallate taste bud cells isolated from *Adipor1*^-/-^ and WT male mice (F (1, 276) = 1.204, P=0.2735), there was a significant interaction (F (4, 276) = 4.160, P=0.0027; Figure 1A). Tukey’s multiple comparisons test revealed that circumvallate taste cells from male *Adipor1*^-/-^ mice showed significantly decreased responsiveness to LA at the highest concentration (100 µM, adjusted P=0.0002; Figure 1A). We did observe a significant genotype effect on LA responses between fungiform taste bud cells isolated from *Adipor1*^-/-^ and WT male mice (F (1, 480) = 5.619, P=0.0182), as well as a significant interaction (F (4, 480) = 2.529, P=0.0399; Figure 1B). Fungiform taste cells from male *Adipor1*^-/-^ mice showed decreased responses to 30 µM (adjusted P=0.0075) and 100 µM LA (adjusted P=0.0023; Figure 1B). In contrast, circumvallate taste bud cells isolated from *Adipor1*^-/-^ females displayed an increase in responsiveness to LA (F (1, 523) = 14.40, P=0.0002; Figure 1C). Tukey’s multiple comparisons test revealed that circumvallate taste cells from female *Adipor1*^-/-^ mice showed significantly increased responsiveness to LA at concentrations of 30 µM (adjusted P<0.0001) and 100 µM (adjusted P=0.0024) (Figure 1C). There was no observed genotype effect (F (1, 359) = 1.343, P=0.2473) and interaction (F (4, 359) = 0.7527, P=0.5567) in LA responsiveness between fungiform taste bud cells isolated from *Adipor1*^-/-^ and WT females (Figure 1D).

**Figure 1.**
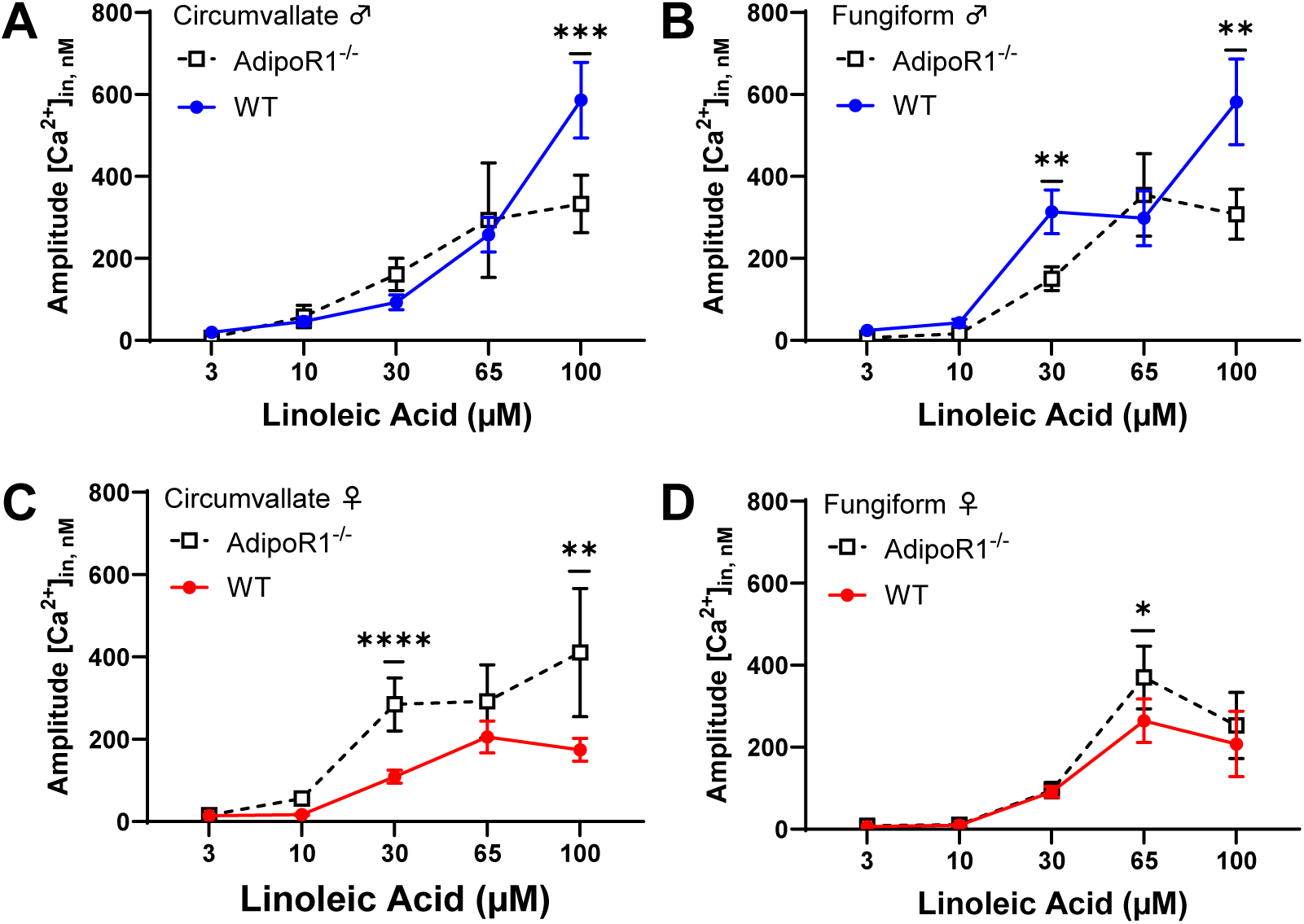
Calcium responses to linoleic acid (LA) in taste cells of *Adipor1*^-/-^ and WT mice. LA (3, 10, 30, 65, and 100 µM) was applied to Fura2-loaded taste bud cells isolated from mice and amplitude measurements were calculated during stimulus response. (A) No significant genotype effect, but a significant interaction (P=0.0027) on LA responses was observed in male circumvallate taste bud cells; (B) Reduced LA responses were observed in *Adipor1*^-/-^ male fungiform taste bud cells (P=0.0182); (C) Increased LA responses were observed in *Adipor1*^-/-^ female circumvallate taste bud cells (P=0.0002); (D) There was no observed genotype effect and interaction on LA responses in female fungiform taste bud cells. Two-way ANOVA was performed to compare the differences between the two genotypes, with Tukey’s multiple comparisons *post hoc* test conducted to determine which LA concentrations differed between the experimental groups. * p < 0.05, ** p < 0.01, *** p < 0.001, **** p < 0.0001.

### Recording of gustatory nerve responses in Adipor1^-/-^ and WT mice

Next, we asked whether global gene knockout of *Adipor1* in mice would have an impact on gustatory nerve responses to various taste stimuli. Whole-nerve recording was performed on chorda tympani (CT) nerves from *Adipor1*^-/-^ and WT mice. Dose-dependent CT nerve responses to different concentrations of intralipid (0.1, 0.5, 1, 2.5, and 5%) were observed in all animal groups, and there was no significant genotype effect on these nerve responses (Figure 2A and 2B). As expected, the nerve responses to sweet (500 mM sucrose), medium-chain saturated fatty acid (0.5 mM, capric acid), bitter (1 mM denatonium benzoate), sour (1 mM HCl), and salty (30 mM NaCl) were similar in the absence or presence of *Adipor1* in both males and females (adjusted P>0.05 for all tests) (Figure 2C and 2D).

**Figure 2.**
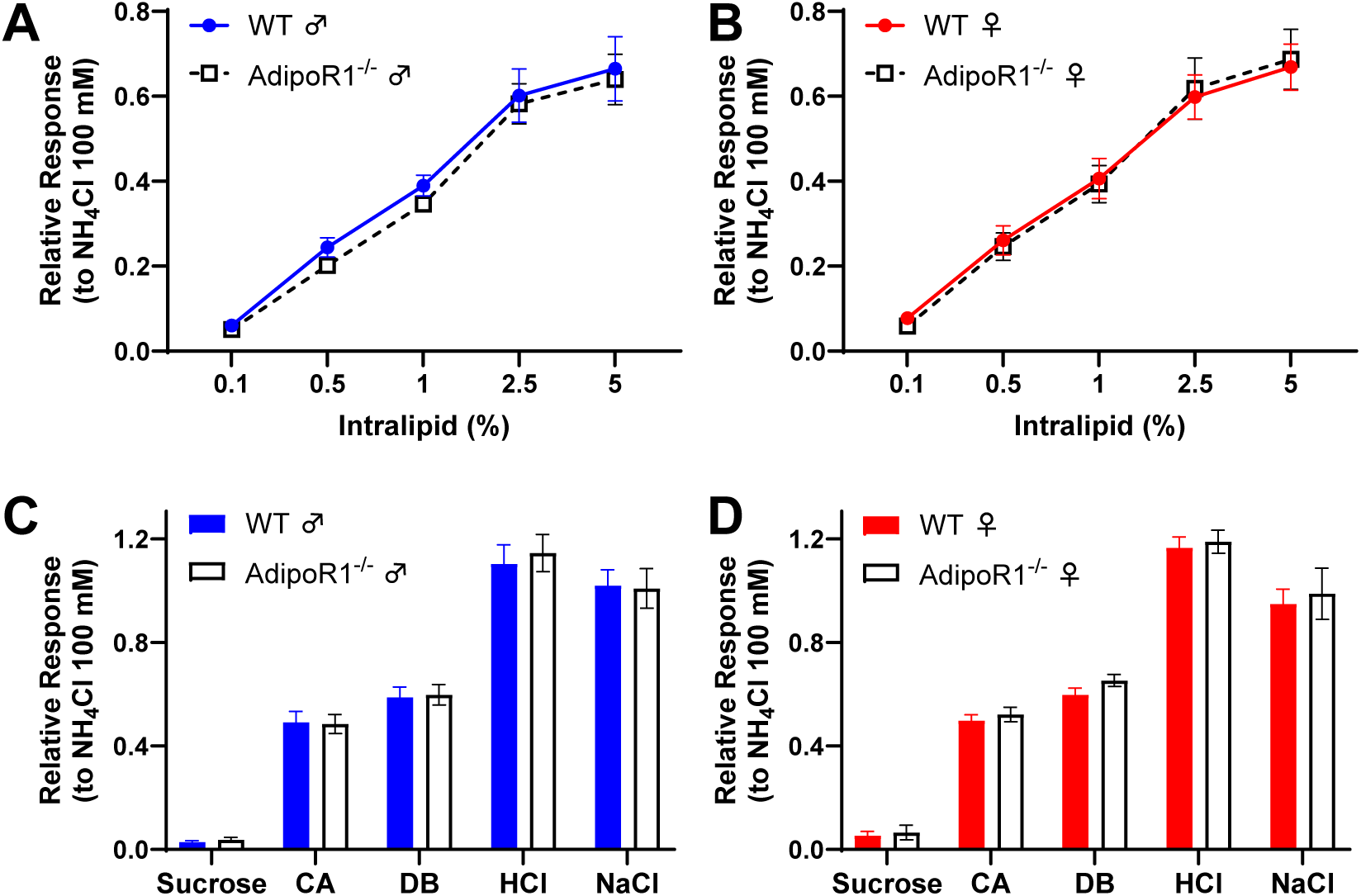
Chorda tympani (CT) nerve responses to different concentrations of intralipid and various tastants (15 s) in *Adipor1*^-/-^ and WT mice. Dose-dependent CT nerve responses to intralipid were observed in all animal groups, and there was no significant genotype effect on these nerve responses in males (A) and females (B); No significant differences were observed between the two genotypes in male (C) and female (D) mice when stimulated with sweet (500 mM sucrose), medium-chain saturated fatty acid (0.5 mM, capric acid, CA), bitter (1 mM denatonium benzoate, DB), sour (1 mM HCl), and salty (30 mM NaCl) tastants. NH4Cl (100 mM) was applied at the beginning and the end of each experiment, and taste responses were normalized to the average NH4Cl response within each session. Area under the curve (AUC) of the nerve response for each tastant was calculated. Summary data show the mean ± SEM (n = 4–7 mice). Two-way ANOVA was performed to compare the differences between the two genotypes, with Tukey’s multiple comparisons *post hoc* test conducted to determine which intralipid concentrations differed. Multiple unpaired t-tests, corrected for multiple comparisons using the Holm-Šídák method, were used to compare the other tastants between the two genotypes.

### Brief-access taste testing of Adipor1^-/-^ and WT mice

To determine the biological function of adiponectin signaling in fat taste behavior, brief-access taste tests were performed to investigate the taste responsiveness to different concentrations of intralipid (0.5, 1, 5, 10, and 20%) in both sexes of *Adipor1*^-/-^ and WT mice. Since we noticed a loss of fat taste in naïve *Adipor1*^-/-^ animals and a recovery of fat taste after exposure to fat (Lin et al., 2024), we compared day 1 and day 5 data of the brief-access taste testing for the two genotypes. On the first day of testing, we observed a significant genotype effect when naïve male mice were sampling the intralipid (F (1, 70) = 21.37, P<0.0001), gene knockout of *Adipor1* in males decreased taste behavioral responsiveness to the 1, 10, and 20% intralipid (adjusted P=0.02, 0.0389, and 0.0013, respectively) (Figure 3A). There was no significant genotype effect (F (1, 70) = 0.008956, P=0.9249) or interaction (F (4, 70) = 0.3236, P=0.8612) observed when naïve female mice were sampling the intralipid (Figure 3B). In contrast, no significant genotype effect (F (1, 70) = 2.222, P=0.1406) and interaction (F (4, 70) = 0.2571, P=0.9044) were observed in male mice on testing day 5 (Figure 3C). However, a significant genotype effect was seen in females on testing day 5 (F (1, 70) = 7.263, P=0.0088) (Figure 3D). Tukey’s multiple comparisons test revealed that *Adipor1*^-/-^ female mice displayed decreased responsiveness to intralipid at the highest concentration (20%, adjusted P=0.0391) (Figure 3D).

**Figure 3.**
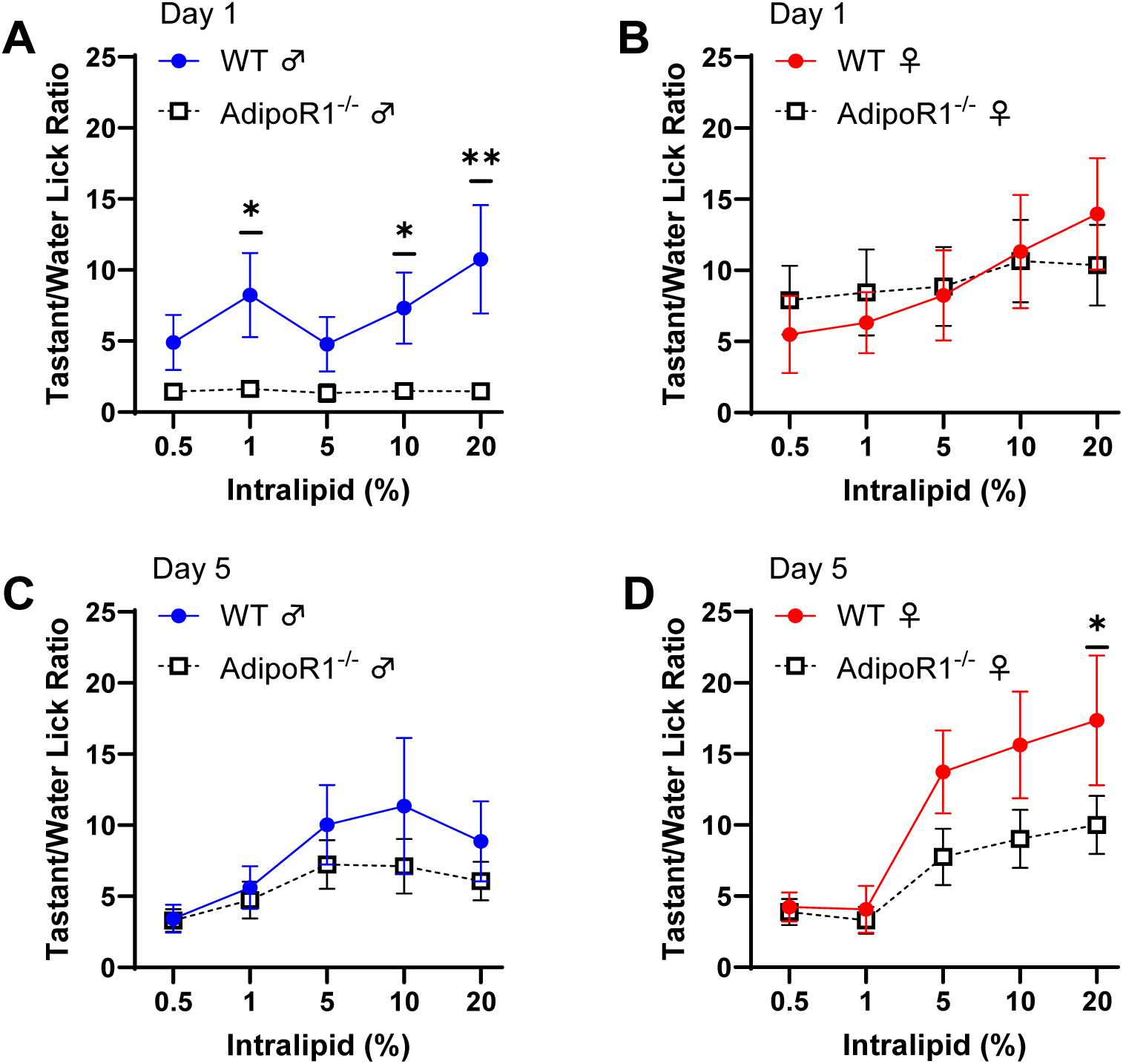
Brief-access taste testing of *Adipor1*^-/-^ and WT mice in response to intralipid. (A) A significant genotype effect was observed when naïve male mice were sampling the intralipid (P<0.0001); (B) No significant genotype effect or interaction was observed when naïve female mice were sampling the intralipid; (C) No significant genotype effect and interaction were observed in fat-experienced male mice on testing day 5; (D) A significant genotype effect was seen in fat-experienced female mice on testing day 5 (P=0.0088). The tastant/water lick ratio was calculated by dividing the mean number of licks per trial for each concentration by the mean number of water licks per trial for that animal. Data are presented as mean ± SEM (n = 8 mice for each group). Two-way ANOVA was performed to compare the differences between the two genotypes, with Tukey’s multiple comparisons *post hoc* test conducted to determine the effect of intralipid concentration. * p < 0.05, ** p < 0.01.

### Adipor1^-/-^ female mice show reduced avoidance to linoleic acid in CTA assays

Next, we sought to determine the gene knockout of *Adipor1* would alter the perceived taste threshold for LA in mice by using conditioned taste aversion (CTA) tests. Animals were injected with LiCl or NaCl (controls) immediately following the oral exposure to the conditioned stimulus (100 μM LA). The results showed that LiCl-injected WT (F (1, 55) = 36.16, P<0.0001) and *Adipor1*^-/-^ (F (1, 60) = 36.76, P<0.0001) male mice developed a similar aversion to LA at concentrations higher than 30 μM LA (Figure 4A and 4B). Furthermore, LiCl-injected WT female mice significantly avoided LA at concentrations as low as 3 μM LA (F (1, 65) = 72.32, P<0.0001) (Figure 4D). In contrast, *Adipor1*^-/-^ female mice did not show any significant aversions for LA with concentrations up to 100 μM LA (F (1, 45) = 0.688, P=0.4112) (Figure 4E). These data suggested that female mice could detect LA at a lower concentration than male mice, and the lack of *Adipor1* in female mice reduced the taste responsiveness to LA. There were no significant aversions to other taste stimuli, including sucrose and NaCl (adjusted P>0.05 for all tests) (Figure 4C and 4F).

**Figure 4.**
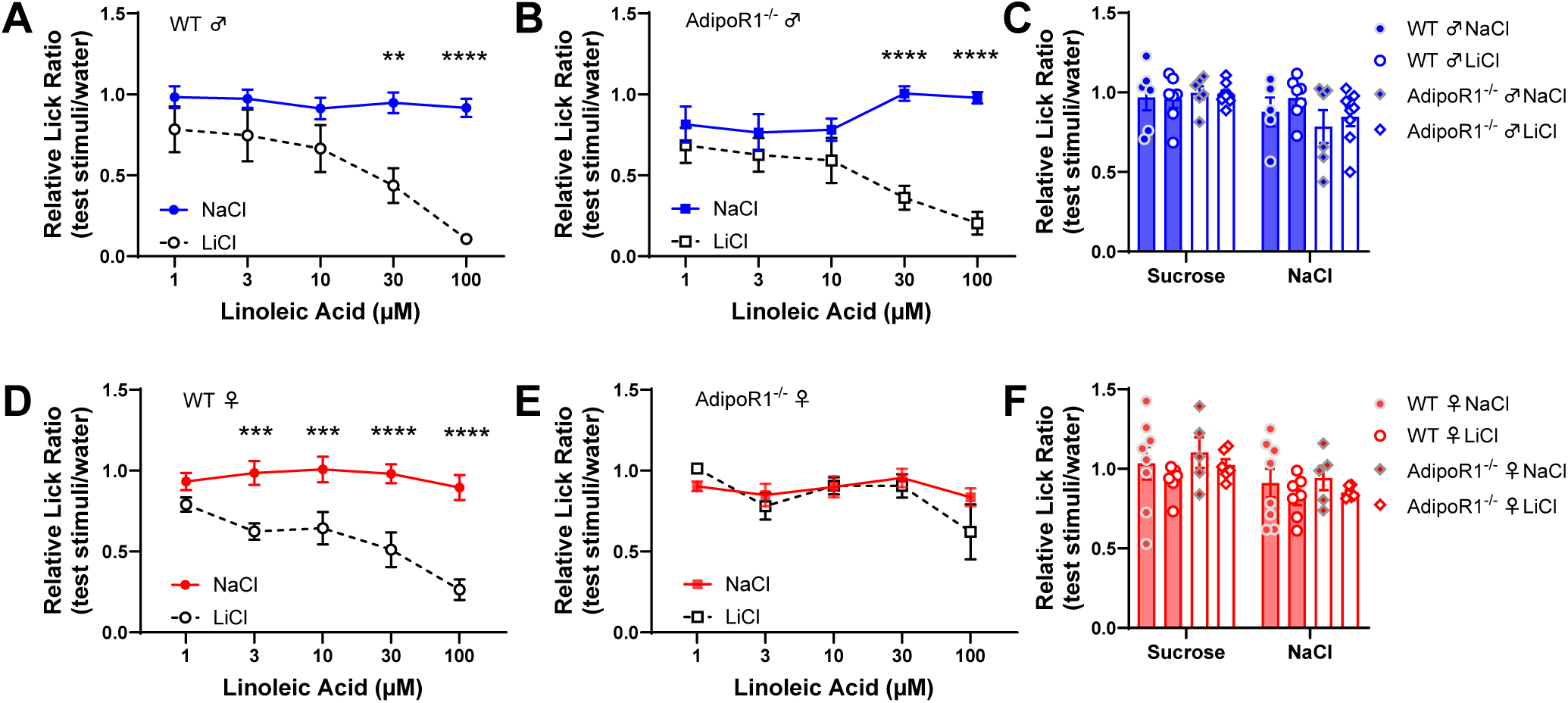
*Adipor1*^-/-^ female mice show reduced avoidance of linoleic acid in the conditioned taste aversion (CTA) assay. (A) LiCl-injected WT (P<0.0001) and (B) *Adipor1*^-/-^ male mice (P<0.0001) developed a similar aversion to LA at concentrations higher than 30 μM LA; (C) No significant aversions to sucrose and NaCl were found in male mice conditioned to avoid fatty acid; (D) LiCl-injected WT female mice significantly avoided LA at concentrations as low as 3 μM LA (P<0.0001); (E) LiCl-injected *Adipor1*^-/-^ female mice did not show any significant aversions for LA with concentrations up to 100 μM LA; (F) No significant aversions to sucrose and NaCl were observed in female mice conditioned to avoid LA. Data are shown as mean ± SEM (n = 5–8 mice). Two-way ANOVA was performed to compare the differences between the two genotypes, with Tukey’s multiple comparisons *post hoc* test conducted to determine which LA concentrations differed. Multiple unpaired t-tests, corrected for multiple comparisons using the Holm-Šídák method, were used to compare the other tastants between the two genotypes. ** p < 0.01, *** p < 0.001, **** p < 0.0001.

### Adipor1^-/-^ female mice show a diminished preference for linoleic acid

To test whether adiponectin signaling impacts fat preference, two-bottle LA preference tests were performed in *Adipor1*^-/-^ and WT mice. In the two-bottle preference assay, LA concentrations ranging from 1 μM to 100 μM LA were plotted for both sexes of *Adipor1*^-/-^ and WT mice. There was no significant difference in preference for LA between male *Adipor1*^-/-^ mice and WT controls (F (1, 70) = 0.0029, P=0.9569) (Figure 5A). In contrast, a significant genotype effect was observed between *Adipor1*^-/-^ and WT female mice (F (1, 60) = 5.107, P=0.0275) (Figure 5B). No difference in preferences for sucrose (sweet), MSG (umami), DB (bitter), and HCl (sour) was observed between the two genotypes in both male and female mice (P>0.05 for all tests) (Figure 5C and 5D). Interestingly, the *Adipor1*^-/-^ mice displayed a significantly higher preference to NaCl (salty) compared to the WT mice in females (t (14) = 3.123, P=0.0075, adjusted P=0.0369), but not in males (t (13) = 1.414, P=0.1807) (Figure 5D).

**Figure 5.**
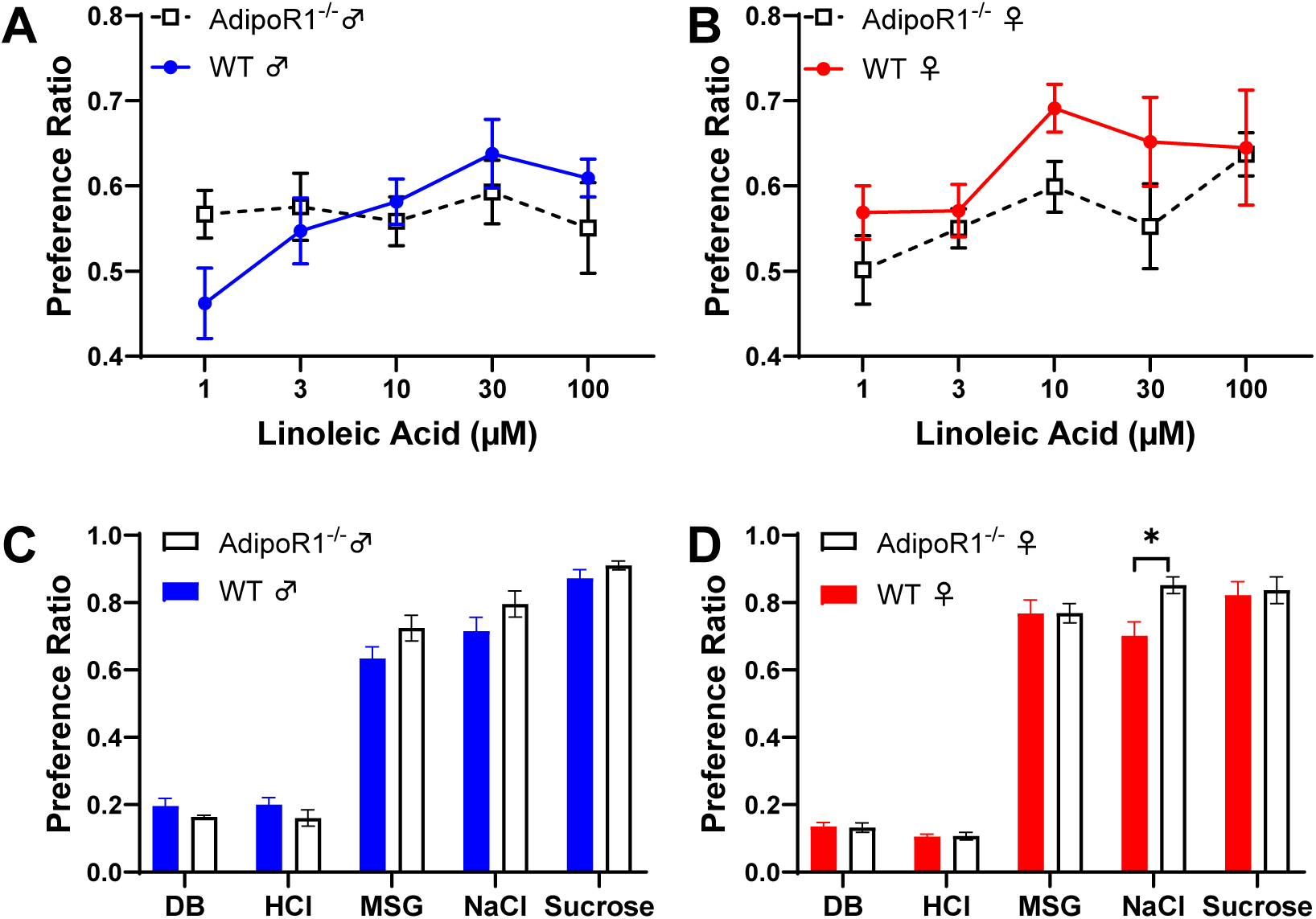
Two-bottle preference test of different concentrations of linoleic acid and various tastants in *Adipor1*^-/-^ and WT mice. (A) There was no significant difference in preference for LA between male *Adipor1*^-/-^ mice and WT controls; (B) A significant genotype effect was observed between *Adipor1*^-/-^ and WT female mice (P=0.0275); (C) No difference in preferences for denatonium benzoate (DB, 1 mM), HCl (1 mM), monosodium glutamate (MSG, 10 mM), NaCl (75 mM), and sucrose (30 mM) was observed between the two genotypes in males; (D) Female *Adipor1*^-/-^ mice displayed a significantly higher preference to 75 mM NaCl (Adjusted P=0.0369) and no difference in preferences for other tastants was observed. Data are shown as mean ± SEM (n = 6–9). Two-way ANOVA was performed to compare the differences between the two genotypes, with Tukey’s multiple comparisons *post hoc* test conducted to determine which LA concentrations differed. Multiple unpaired t-tests, corrected for multiple comparisons using the Holm-Šídák method, were used to compare the other tastants between the two genotypes. * p < 0.05.

### Adipor1^-/-^ and WT mice display similar preferences and intakes for intralipid

Preference for intralipid was then tested using the two-bottle preference assay. This test compared the intralipid preferences and intakes of *Adipor1*^-/-^ and WT mice over a range of intralipid concentrations from 0.1% to 5%. Intralipid intake simply increased with increasing concentration in all animal groups. Although *Adipor1*^-/-^ males consumed more intralipid than WT males at the 2.5% concentration (Tukey’s multiple comparisons test, adjusted P=0.0116), no significant genotype effects were observed in intralipid consumption in both males (F (1, 66) = 2.888, P=0.094) and females (F (1, 60) = 0.3337, P=0.5656) (Figure 6A and 6B). The animals displayed strong preferences for intralipid at 0.5% and higher concentrations, and there was no significant difference in intralipid preference between *Adipor1*^-/-^ and WT mice (males: F (1, 66) = 1.795, P=0.1849; females: F (1, 60) = 0.0001, P=0.9908) (Figure 6C and 6D). Possibly due to the loss of fat taste, naïve *Adipor1*^-/-^ males trended toward having a reduced preference for intralipid at concentrations of 0.1% (adjusted P=0.0568) and 0.5% (adjusted P=0.0678) (Figure 6C).

**Figure 6.**
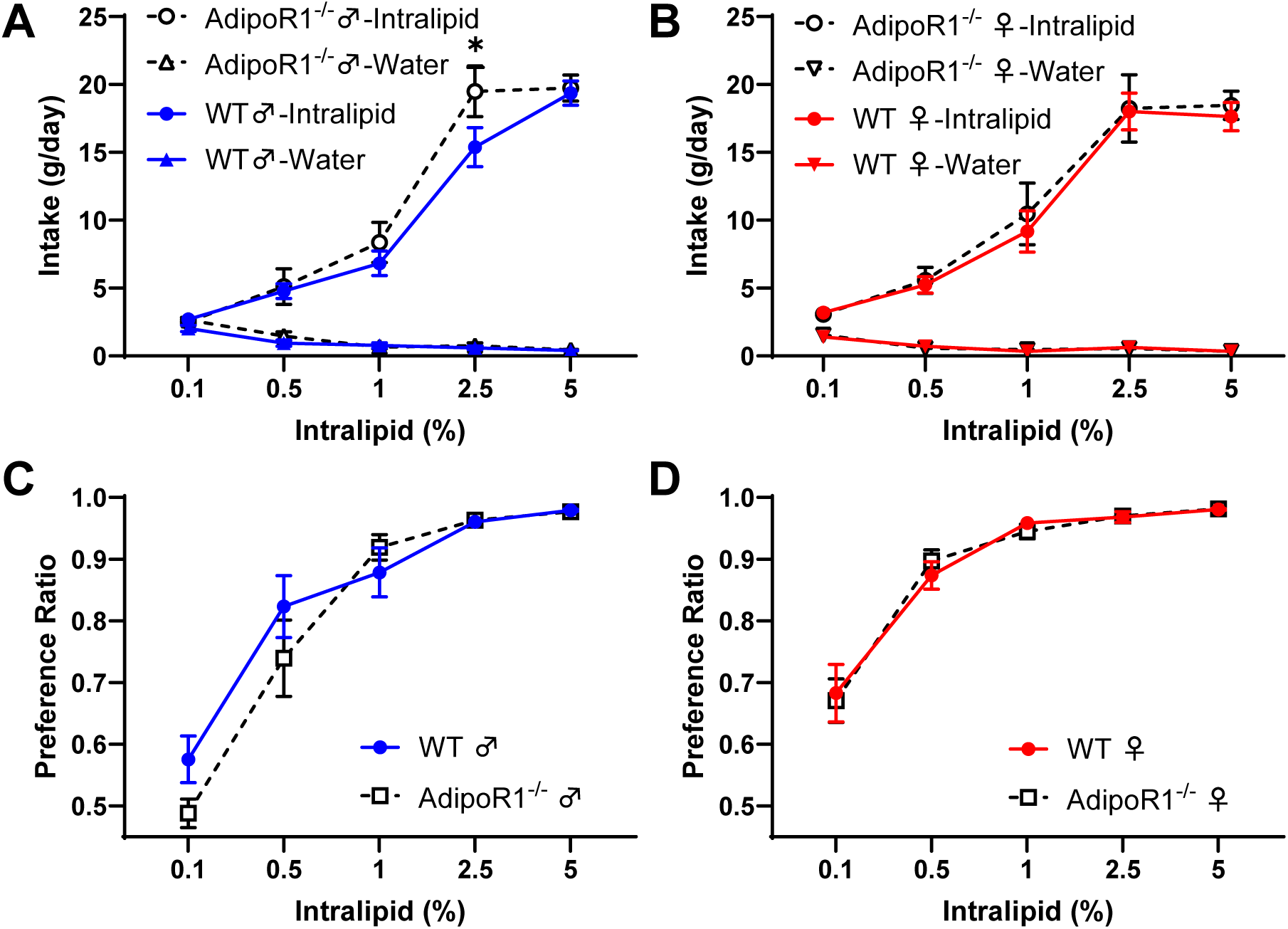
The intake and preference of different concentrations of intralipid in *Adipor1*^-/-^ and WT mice. No significant genotype effect and interaction were observed in intralipid consumption in both males (A) and females (B); There was no significant difference in intralipid preference between *Adipor1*^-/-^ and WT mice in both males (C) and females (D). The ratio of 48-hour tastant intake over the total fluid (tastant + water) consumption was calculated as the preference ratio of the tastant. Data are presented as mean ± SEM (n = 7–8). Two-way ANOVA was performed to compare the differences between the two genotypes, with Tukey’s multiple comparisons *post hoc* test conducted to determine which intralipid concentrations differed. * p < 0.05.

### Adipor1^-/-^ and WT mice display a very similar preference and intake for high-fat diet

To study the effects of adiponectin signaling on food choice, we therefore tested high-fat diet palatability in *Adipor1*^-/-^ and WT mice. Similar to the two-bottle preference assay, singly housed animals were given simultaneous free access to a water bottle, and two hoppers containing control diet (10% calories from fat and 70% calories from carbohydrate) and high-fat diet (60% calories from fat and 20% calories from carbohydrate) in the Promethion system. Not surprisingly, the animals consumed more high-fat diet than the control diet in both males (F (1, 26) = 285.9, P<0.0001) and females (F (1, 28) = 120.5, P<0.0001) (Figure 7A and 7D). The *Adipor1*^-/-^ and WT mice did not significantly differ in their intakes and preferences for high-fat diet (P>0.05 for all tests) (Figure 7A, 7B, 7D and 7E). A significantly higher respiratory exchange ratio was observed in *Adipor1*^-/-^ animals (males: t (14) = 2.711, P=0.01688, adjusted P=0.0335; females: t (14) = 3.454, P=0.00387, adjusted P=0.0077) during the dark period on a high-fat diet (Figure 7C and 7F).

**Figure 7.**
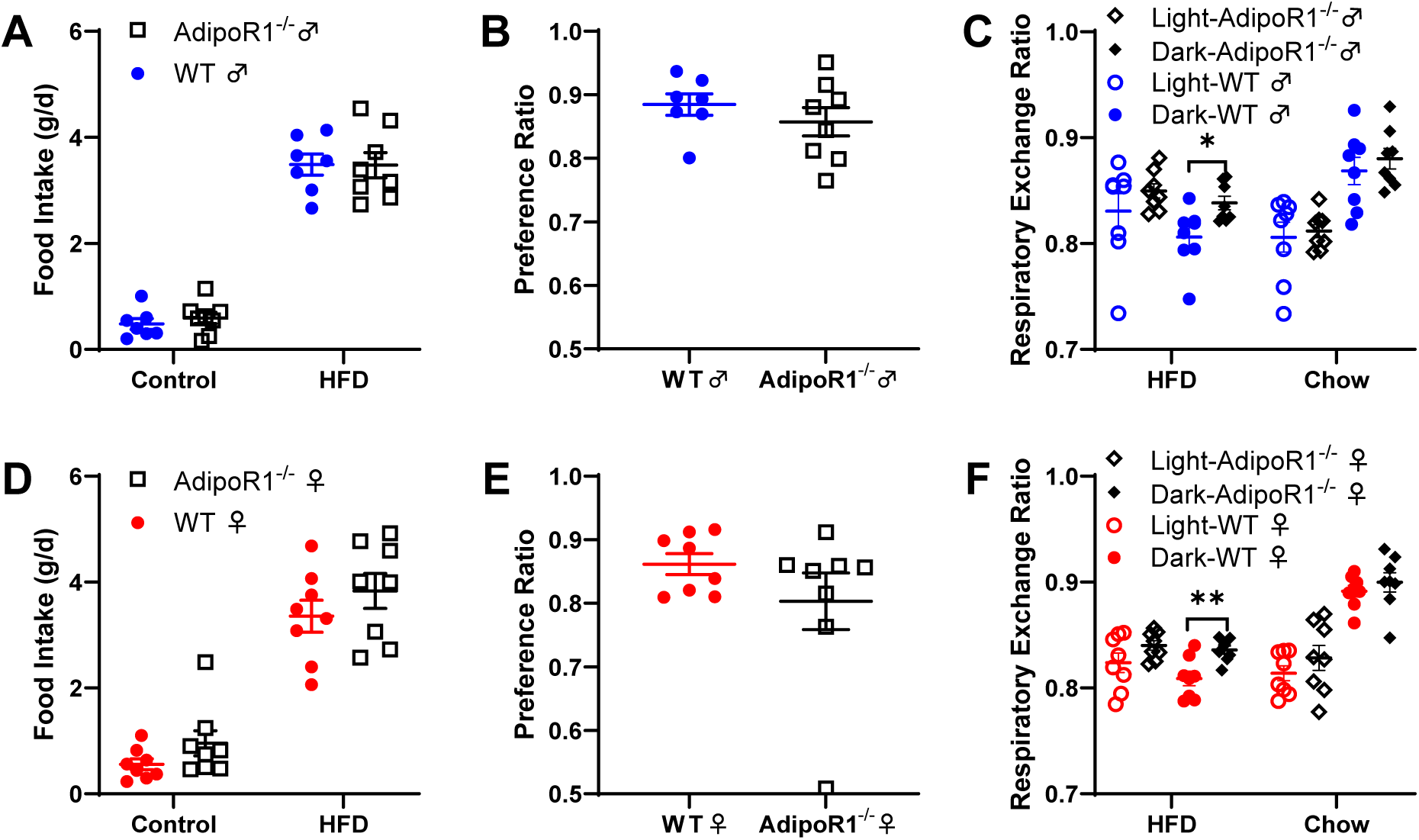
Food intake and preference of *Adipor1*^-/-^ and WT mice with free access to control and high-fat diets. (A) Male mice consumed more high-fat diet than the control diet (P<0.0001), but did not differ in their food intakes between the two genotypes; (B) *Adipor1*^-/-^ and WT males did not significantly differ in their preference for high-fat diet; (C) A significant higher respiratory exchange ratio (RER) was observed in *Adipor1*^-/-^ males during the dark period on a high-fat diet (adjusted P=0.0335); (D) Female mice consumed more high-fat diet than the control diet (P<0.0001), but not differ in their food intakes between the two genotypes; (E) *Adipor1*^-/-^ and WT females did not significantly differ in their preference for high-fat diet; (F) Higher RER was observed in *Adipor1*^-/-^ females during the dark period on a high-fat diet (Adjusted P=0.0077). The preference ratio for the high-fat diet was calculated as high-fat diet intake over total food consumption. Data are presented as mean ± SEM (n = 8 for each group). Two-way ANOVA was performed to compare the differences in food intakes between the two genotypes. An unpaired t-test was performed for the diet preference between the two genotypes. Multiple unpaired t-tests, corrected for multiple comparisons using the Holm-Šídák method, were used to compare the RER between the two genotypes. * p < 0.05, ** p < 0.01.

### Body weight and dietary intakes in HFD fed Adipor1^-/-^ and WT mice

It has been suggested that a high-fat diet and diet-induced obesity modify taste sensitivity (Harnischfeger & Dando, 2021; Maliphol et al., 2013). To test whether adiponectin signaling was involved in the influence of diet-induced taste changes, animals were given free access to a 60% high-fat diet. The feeding study was stopped when the animals were under a high-fat diet for 5 (males) and 4 (females) weeks. Male animals showed no significant differences in body weight gain (F (1, 70) = 0.4701, P=0.4952) (Figure 8A), and no significant changes in free fluid and fat mass were found between *Adipor1*^-/-^ and WT males (P>0.05 for all tests), but WT males gain more lean mass compared to *Adipor1*^-/-^ males (t (14) = 3.201, P=0.0064, adjusted P=0.0191) (Figure 8B). Moreover, no significant differences in food and water consumptions were observed between *Adipor1*^-/-^ and WT males (P>0.05 for all tests) (Figure 8C). In contrast, a significant genotype effect was observed in female animals under a high-fat diet (F (1, 60) = 6.016, P=0.0171), *Adipor1*^-/-^ females trended to gain more body weights than WT controls (Figure 8D), however no significant changes were found between *Adipor1*^-/-^ and WT females in free fluid, lean, and fat mass (P>0.05 for all tests) (Figure 8E). Surprisingly, while they trended to gain more body weight, *Adipor1*^-/-^ females seemed to consume less food and water compared to the WT controls, but not significantly so (food: t (12) = 1.806, P=0.096; water: t (12) = 1.598, P=0.136) (Figure 8F). An increased body weight gain was observed in the male but not female, *Adipor1*^-/-^ mice, with reduced energy expenditure and spontaneous locomotor activity in a previous study (Bjursell et al., 2007). However, opposite findings were observed in our experiments between the two genders. Similar to the body weight gain, no significant differences in energy expenditure and pedestrian locomotion were found between *Adipor1*^-/-^ and WT males (Figures S1A, S1B, S2A, S2B; *Supplement*). In contrast, female *Adipor1*^-/-^ mice showed reduced energy expenditure and pedestrian locomotion, especially under a high-fat diet (Figures S1C, S1D, S2C, S2D; *Supplement*).

**Figure 8.**
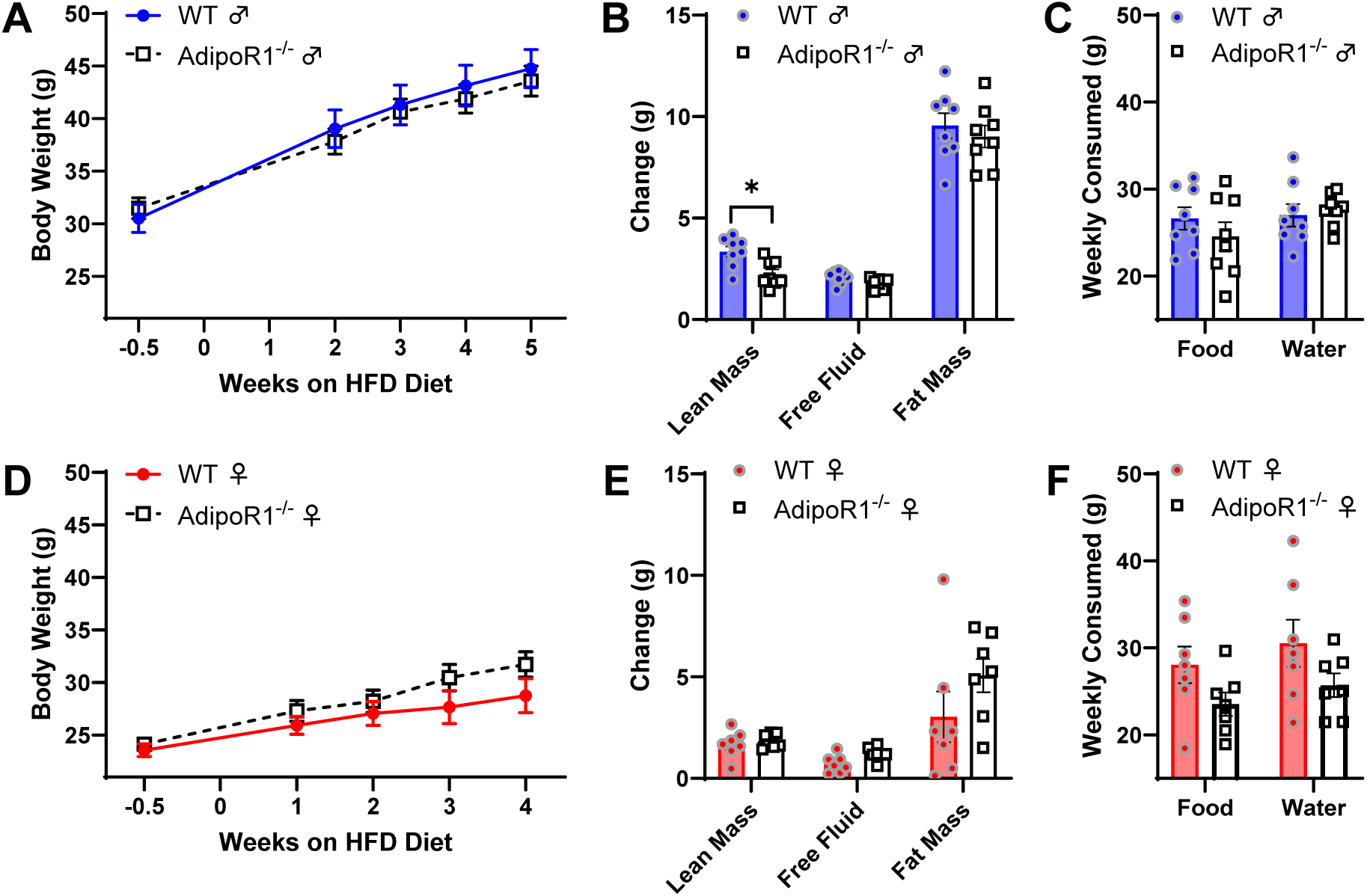
Body weight, body composition change, food and water intake in *Adipor1*^-/-^ and WT mice on a high-fat diet. (A) Male animals showed no significant differences in body weight gain under a high-fat diet; (B) WT males gain more lean mass on 5 weeks of high-fat diet compared to *Adipor1*^-/-^ males (adjusted P=0.0191), and no significant changes in free fluid and fat mass were found between *Adipor1*^-/-^ and WT males; (C) No significant differences in food and water consumptions were observed between *Adipor1*^-/-^ and WT males; (D) A significant genotype effect on body weight gain was observed in female animals under a high-fat diet (P=0.0171); (E) No significant changes in free fluid, lean, and fat mass were found between *Adipor1*^-/-^ and WT females; (F) No significant differences in food and water consumptions were observed between *Adipor1*^-/-^ and WT females. Data are presented as mean ± SEM (n = 7–8 mice). Two-way ANOVA was performed to compare the differences between the two genotypes, with Tukey’s multiple comparisons *post hoc* test conducted to determine if the body weight differed. Multiple unpaired t-tests, corrected for multiple comparisons using the Holm-Šídák method, were used to compare the body composition change, food, and water intake between the two genotypes. * p < 0.05.

### Neural and behavioral taste responses in HFD fed Adipor1^-/-^ and WT mice

High-fat diet fed animals received chow diet for several days, and we then subjected these animals to a two-bottle preference test for a series of intralipid concentrations, followed by CT nerve recordings to investigate whether there were any changes in fat taste responsiveness. Similarly, the animals did not differ in their intralipid intakes and preferences between *Adipor1*^-/-^ and WT mice (P>0.05 for all tests) (Figure S3A, S3B; *Supplement*). All animals displayed strong preferences (>90%) for intralipid at 0.1% and higher concentrations (Figure S3C, S3D; *Supplement*). In addition, no significant genotype effect was seen between high-fat fed *Adipor1*^-/-^ and WT animals for the CT nerve responses to intralipid (Figure S4A, S4B; *Supplement*). Except for the capric acid responses in males (t (9) = 3.682, P=0.005, adjusted P=0.025), neural responses of high-fat fed *Adipor1*^-/-^ mice did not differ from their WT controls for any other taste stimuli in both genders (adjusted P>0.05 for all tests) (Figure S4C, S4D; *Supplement*).

## Discussion

As a major component of many common foods and one of the three main macronutrients in human and animal diets, the biological importance of fats has been well documented (Calder, 2015; Kumar et al., 2017). Fats provide most living beings with an important source of dense energy and essential fatty acids, facilitate the absorption of fat-soluble micronutrients, and participate in critical metabolic and structural functions (Bhattacharya & Rattan, 2021). Energy-dense foods rich in fats are especially good tasting and taste is one of the primary driving forces of dietary choice and consumption (Drewnowski, 1997; Mizushige et al., 2007). Therefore, high-fat foods are generally preferred in humans and animals (Drewnowski & Greenwood, 1983; Smith et al., 2000). However, excessive dietary fat intake has been linked to increased risk of several serious health conditions, including obesity (Bray & Popkin, 1998), type II diabetes (Salmeron et al., 2001), cardiovascular disease (Zeng et al., 2023), and even certain types of cancer (Sieri et al., 2008; Zhao et al., 2016). Increasing evidence suggests that fat taste responsiveness could be regulated by several hormones, and this hormonal modulation of fat taste may influence fat preference and dietary fat intake (Calder et al., 2021; Crosson et al., 2019; Martin et al., 2012; Ullah et al., 2021). In the present study, we observed that disruption of adiponectin signaling through global genetic knockout of *Adipor1* in mice altered the animals’ cellular and behavioral taste responsiveness to fatty acids in a sex-dependent manner, however, little or no effect on dietary fat intake was found.

To date, adiponectin (Crosson et al., 2019; Lin et al., 2023), cannabinoids/cannabinoid 1 receptor (CB1R) (Brissard et al., 2018), ghrelin/ghrelin receptor (GHS-R) (Cai et al., 2013; Calder et al., 2021), glucagon-like peptide-1 (GLP-1)/GLP-1R (Martin et al., 2012; Treesukosol & Moran, 2022), leptin/leptin receptor (Ob-Rb) (Kitagawa et al., 2007; Ullah et al., 2021), and peptide YY (PYY) (La Sala et al., 2013) are implicated in the modulation of taste responsiveness to fatty acids. Endocannabinoids (Niki et al., 2015; Yoshida et al., 2010), GLP-1 (Martin et al., 2009; Shin et al., 2008), and leptin (Kawai et al., 2000; Niki et al., 2015; Shigemura et al., 2004; Yoshida et al., 2021; Yoshida et al., 2015) are also well-documented sweet taste modulators. Leptin suppresses mouse taste cell responses and preference to fatty acid and these effects were mediated by Ob-Rb on taste bud cells (Ullah et al., 2021). In addition, CD36, leptin, and Ob-Rb are co-expressed in mouse taste bud cells and leptin signaling interacts with CD36 mediated fat taste transduction (Ullah et al., 2021). CD36 protein levels in mouse taste cells have been shown to be downregulated by fat in the diet, and that change in CD36 levels in taste cells appears to be sufficient to modulate the motivation for fat during a meal (Martin et al., 2011). Although there were no significant differences in CD36 gene and protein levels between taste cells from WT and *Glp1r*^-/-^ mice, the postprandial downregulation of CD36 protein levels found in WT mouse taste cells was not observed in taste bud cells of refed *Glp1r*^-/-^ animals (Martin et al., 2012). GLP-1 signaling is also involved in the modulation of fat taste and *Glp1r*^-/-^ male mice were unable to taste low concentrations of oil compared to the controls (Martin et al., 2012). A reduced taste cell responses and preference to fatty acids have also been found in *Cb1r*^-/-^ male mice (Brissard et al., 2018). No significant changes in CD36 protein levels were seen in taste cells of *Cb1r*^-/-^ mice compared with WT animals, however, a low GLP-1 basal level and a decreased *Glp1r* expression level were exhibited in *Cb1r*^-/-^ mice (Brissard et al., 2018).

Additionally, a reduced GPR120 expression has been found in isolated taste bud cells from ghrelin-deficient mice, suggesting that ghrelin signaling and GPR120 may interact to modulate fat taste (Cai et al., 2013). Recently, it has been reported that CD36 protein level was significantly decreased in the retina of *Adipor1*^-/-^ mice (Lewandowski et al., 2024). Further studies are necessary to examine the gene expression and protein levels of CD36 in the taste buds of *Adipor1*^-/-^ mice. The present study found sex-dependent changes in cellular and behavioral taste responses to fatty acids in *Adipor1*^-/-^ mice. Our findings indicate that adiponectin may selectively enhance fat taste sensitivity in mice via *Adipor1* on the taste cells. Reduced taste responses to fatty acids were found in *Adipor1*^-/-^, *Cb1r^-/-^*, *Glp1r^-/-^*, and *Ghsr^-/^*^-^ animals. Whether these signaling pathways interact to influence fat taste remains an open question. However, their modulation has different targets [CD36 (adiponectin/AdipoR1) vs. GPR120 (ghrelin/GHS-R) vs. CD36 and GPR120 (GLP-1/GLP-1R)] and taste specificities [fat (adiponectin/AdipoR1) vs. fat-sweet (endocannabinoids/CB1R) vs. fat-sweet-sour (GLP-1/GLP-1R) vs. fat-sweet-sour-salty (ghrelin/GHS-R)], suggesting that these hormones, as well as mediated signaling pathways may play different roles in the modulating of fat taste.

AdipoR1 and AdipoR2 are the two main functional receptors for adiponectin mediating lipid and glucose metabolism, with AdipoR1 activating the AMPK signaling and AdipoR2 activating the peroxisome proliferator-activated receptor α (PPARα) signaling (Yamauchi et al., 2007). It has been proposed that AdipoR1 and AdipoR2 are the *yin-yang* receptors of adiponectin in energy metabolism, as mice genetically deficient in AdipoR1 and AdipoR2 have remarkably opposite phenotypes, producing opposite effects on energy expenditure and physical activity (Bjursell et al., 2007). Despite similar food intake to WT controls, an increased body weight gain over time was observed in male but not female *Adipor1*^-/-^ mice in a previous study (Bjursell et al., 2007). In our study, no differences were found in body weight and body weight gain in *Adipor1*^-/-^ males before and after the high-fat diet feeding study, whereas *Adipor1*^-/-^ females gained more weight under a high-fat diet, and this was not due to increased fat intake.

A significantly enhanced reduction in respiratory exchange ratio was found in high-fat diet-fed WT animals compared with *Adipor1*^-/-^ mice, suggesting a greater ability to utilize dietary fat as a fuel source in WT mice. However, this was found in both genders and cannot explain the sex-dependent body weight change. Differences in age and dietary status in these two studies may account for the sex-specific differential regulation of body weight by adiponectin signaling. Nonetheless, the causes of body weight gain over time in *Adipor1*^-/-^ males under chow diet and *Adipor1*^-/-^ females fed a high-fat diet appear to be the same, with reduced energy expenditure and spontaneous locomotor activity in both. The obvious changes in animals’ pedestrian locomotion were seen during the transition from chow to high-fat diet feeding. Thus, it is difficult to exclude the implication between taste signals and the subsequent series of metabolic changes. These studies suggest that adiponectin/AdipoR1 signaling plays a crucial role in maintaining healthy body weight, primarily by regulating the expenditure side of the energy balance equation rather than through dietary intake.

Under the chow diet, the role of adiponectin/AdipoR1 signaling in female body weight control may be less important, or there may be a complementary mechanism in females lacking *Adipor1*. However, targeting AdipoR1 may help develop therapeutic strategies for obesity in aging males caused by disruption of adiponectin signaling. Evidence of adiponectin resistance has been shown in high-fat diet feeding animals (Mullen et al., 2009; Mullen et al., 2007). High-fat diet feeding may cause adiponectin resistance in WT male mice, which results in no significant difference in body weight gain regardless of *Adipor1* deficiency. Estradiol has been shown to play an important role in the modulation of adiponectin sensitivity and could overcome adiponectin resistance in diabetic mice by upregulating AdipoR1 levels in the skeletal muscle (Chattopadhyay et al., 2022). Additionally, increased AdipoR1 protein levels were found in certain tissues of *Adipor2^-/-^* mice, and these animals were resistant to high-fat diet-induced obesity (Bjursell et al., 2007). These findings support the importance of adiponectin/AdipoR1 signaling in body weight control and may explain why the WT females are more resistant to the obesogenic effects of high dietary fat. Therefore, targeting adiponectin/AdipoR1 signaling may help discover effective ways to alleviate high-fat diet-induced obesity.

Three commonly used taste behavioral assays (two-bottle preference tests, brief-access taste tests, and conditioned taste aversion tests (Gaillard & Stratford, 2016) were used to determine if and how global genetic knockout of *Adipor1* affects taste-related behaviors in mice. The two-bottle preference assay assesses whether fatty acids are preferred over water and measures broad taste preferences and provides clues to relative senstivities. This assay was favored for its simplicity and ease of implementation, but it is the most subject to post-ingestive influences (Ackroff & Sclafani, 2014). The brief-access taste assay was then introduced to minimize the post-oral effects and increase our confidence that the observed behavior differences between groups, if any, arise solely from peripheral gustatory signals. Conditioned taste aversion assay was applied to determine the specific taste threshold for fatty acids. Similar to body weight control, adiponectin/AdipoR1 signaling also modulates fat taste in sex-specific ways. In males, we found reduced LA-induced calcium responses in taste cells isolated from *Adipor1*^-/-^ mice and a taste loss in brief-access taste test when *Adipor1*^-/-^ animals initially exposed to intralipid. Interestingly, the fat taste loss in fat-naïve *Adipor1*^-/-^ males was restored after two days of exposure to intralipid and showed no significant difference with WT controls. This experience-induced fat taste recovery may explain why no differences in taste behavior between the two genotypes were found in two-bottle preference (naïve *Adipor1*^-/-^ male showed no preference for 0.1% intralipid and slightly lower preference for 0.5% intralipid compared to WT mice, then similar preference for 1, 2.5, and 5% intralipid) and conditioned taste aversion tests (fat taste may have recovered from the three consecutive days of fatty acid exposure on condition days prior to testing).

In females, we observed a surprising increase in LA-induced calcium responses in taste cells isolated from *Adipor1*^-/-^ mice compared with WT controls, suggesting that naïve *Adipor1*^-/-^ females may have developed an unknown complementary mechanism to assist in sensing fatty acids. Interestingly, *Adipor1*^-/-^ females were found to have reduced gustatory behavioral responses to fatty acids in all three behavioral assays (diminished preference for LA in two-bottle preference test, lower intralipid/water lick ratio in brief-access assay, and reduced avoidance for LA in conditioned taste aversion assay). Although there is limited literature discussing the role of adiponectin signaling in the taste system, a recent study demonstrated that sex differences are linked to human olfactory sensitivity via adiponectin levels (Pfabigan et al., 2022). Similar sex-specific hormonal modulation of fat taste responsiveness was also found in ghrelin receptor *Ghsr* knockout mice (Calder et al., 2021). Our results further support previous findings that there are significant differences in fat taste thresholds between WT males and females (Dahir et al., 2021; Pittman et al., 2008).

The mechanisms responsible for these sex-specific differences remain unknown; however, gender differences in fat taste are not uncommon, and estradiol is a primary modulator that alters fatty acid taste responsiveness (Dahir et al., 2021). Evidence indicates that leptin resistance developed when the rats consumed carbohydrate-containing solutions rather than calorie-free sweet solutions, suggesting that the development of leptin resistance in rats is not associated with sweet taste (Harris, 2019). Adiponectin resistance has been demonstrated in animals fed high-fat diets (Mullen et al., 2009; Mullen et al., 2007). However, it remains unclear whether the development of adiponectin resistance is related to fat taste. In our taste behavioral studies, we noticed that there was no difference in taste behavioral responses to fatty acids between fat-experienced *Adipor1*^-/-^ and WT males, suggesting adiponectin/AdipoR1 signaling may not be required for the fat taste modulation in fat-experienced males. In other words, we speculate that the taste system may rapidly develop resistance to adiponectin after exposure to fat. Previous studies have shown that globular adiponectin fails to increase fatty acid oxidation in skeletal muscle of rats fed a high-fat diet, which occurs extremely rapidly (less than 3 days) (Mullen et al., 2009). Diabetic mice have been reported to develop sex-specific (male but not female) resistance to globular adiponectin treatment, and estradiol could impart adiponectin sensitivity via its nuclear receptors (Chattopadhyay et al., 2022). Therefore, we speculate that the sex-specific modulation of fat taste behavior in mice by genetic knockout of *Adipor1* may indeed be the case.

Control of food choice and intake is complex, with numerous factors influencing food selection and dietary intake. Among them, taste has been considered by consumers to be one of the primary determinants of food choice. Many hormones and peptides involved in the regulation of food intake also play a key role in the modulation of taste, such as neuropeptide Y (NPY), PYY, ghrelin, GLP-1, leptin, cholecystokinin (CCK), and adiponectin. In the present study, the demonstration that no differences in dietary fat preference and intake were found between WT and *Adipor1*^-/-^ mice, suggesting that adiponectin/AdipoR1 signaling is either not essential for the control of dietary fat intake or multiple compensatory mechanisms exist for the modulation of dietary fat choice and intake. Indeed, dietary fat intake has been shown to be selectively reduced by enterostatin (Okada et al., 1991) and increased by galanin (Karatayev et al., 2009). It has also been reported that NPY (Primeaux et al., 2006), PYY (Dischinger et al., 2019), GLP-1 (Dischinger et al., 2019), ghrelin (Shimbara et al., 2004), amylin (Mack et al., 2007), and proopiomelanocortin (POMC) (Tung et al., 2007) could influence dietary fat choice, preference, and intake. Further research is needed to explore the link between fat taste and dietary fat selection and intake.

## Materials and Methods

### Animals

The mouse strain (B6.129P2-*Adipor1*^tm1Dgen^/Mmnc, RRID: MMRRC_011599-UNC) was donated to the NIH-sponsored Mutant Mouse Resource and Research Center (MMRRC) by Deltagen. Heterozygous *Adipor1*^+/-^ mice were ordered from the MMRRC facility at the University of North Carolina. When the offspring mice produced by heterozygous intermating were genotyped, new homozygous breeding was established to produce a larger amount of age-matched *Adipor1*^-/-^ mice and *Adipor1*^+/+^ controls for experiments. Mice were housed under standard laboratory conditions (12 h:12 h day/night cycle) with water and normal chow available *ad libitum* unless otherwise specified. All animal procedures were approved by the Institutional Animal Care and Use Committee of the University of Central Florida.

### Diets and solutions

The high-fat diet (D06062303) and control diet (D07020902) were ordered from Research Diets (New Brunswick, NJ, USA). High fat diet consisted of 60% fat (with 3.3:1 unsaturated fatty acids to saturated fatty acids sourced mostly from lard and safflower oil), 20% carbohydrates, and 20% protein with an energy density of 5.24 kcal/gram, while the control diet consisted of 10% fat (with equal unsaturated to saturated fatty acids ratio), 70% carbohydrates (primarily high corn starch), and 20% protein with an energy density of 3.85 kcal/gram.

Standard Tyrode’s solution contained (in mM) 140 NaCl, 5 KCl, 1 CaCl_2_, 1 MgCl_2_, 10 HEPES, 10 glucose, and 10 Na pyruvate; adjusting the pH to 7.40 with NaOH; 300–320 mOsm. Calcium-magnesium free Tyrode’s solution contained (in mM): 140 NaCl, 5 KCl, 2 BAPTA, 10 HEPES, 10 glucose, and 10 Na pyruvate; adjusting the pH to 7.40 with NaOH; 300–320 mOsm. Linoleic acid (LA) was purchased from Sigma (St. Louis, MO, USA), prepared as stock solutions (25 mg/ml) in 100% ethanol, and then stored under nitrogen at −20 °C. LA working solutions (1–100 μM) were made from stock solutions immediately before use. Intralipid 20% IV fat emulsion was purchased from Patterson Veterinary (Loveland, CO, USA) and diluted with pure water to produce the concentrations required in each experiment. Enzyme cocktail components collagenase A, dispase II, and trypsin inhibitor were obtained from Sigma.

### Taste bud cell isolation

Individual taste bud cells were isolated from mouse circumvallate papilla and fungiform papilla following procedures used in previous reports (Dahir et al., 2021; Liu et al., 2021). Briefly, adult males and females from both *Adipor1^-/-^* mice and *AdiporR1^+/+^* controls were sacrificed by exposure to CO_2_ and followed by cervical dislocation. Tongues were removed and immediately placed in Tyrode’s solution. The anterior portion of the tongue containing the fungiform papilla and the area surrounding the circumvallate papilla were then injected with an enzyme cocktail in Tyrode’s solution [collagenase A (0.5 mg/ml), dispase II (2 mg/ml), and trypsin inhibitor (1 mg/ml)]. The injected tongue was incubated in Tyrode’s solution and bubbled with O_2_ for approximately 40 minutes. The lingual epithelium was peeled from the underlying muscle layer and pinned flat in Tyrode’s solution in a Sylgard-coated petri dish. Next, the epithelium was incubated in calcium-magnesium-free Tyrode’s solution for 5–7 minutes in males or 3–5 minutes in females, washed with standard Tyrode’s solution, and subsequently placed in the same enzyme cocktail described above for another 3–5 minutes (males) or 1–3 minutes (females). Taste bud cells were isolated by gentle suction using a fire-polished glass pipette under a dissection microscope and placed on coverslips coated with Corning Cell-Tak Cell and Tissue Adhesive (Corning, PN, USA) for calcium imaging.

### Calcium imaging

Intracellular calcium imaging was carried out on isolated taste bud cells loaded with 4 µM of the ratiometric calcium indicator Fura-2-acetoxymethyl ester (Fura-2 AM, Invitrogen) in Tyrode’s with 0.05% pluronic acid F-127 (Invitrogen) for about 1 hour at room temperature in the dark. The coverslips with taste bud cells ready for imaging were placed onto a perfusion chamber (RC-25F, Warner Instruments, Holliston, MA, USA). A series of LA solutions (1–100 μM) were applied by a bath perfusion system at a flow rate of 4 mL/minute for 3 minutes, followed by 1 minute of 0.1% fatty acid-free BSA solution, and then regular Tyrode’s for about 2 minutes to remove the LA until the calcium signal returned to near baseline level. Taste bud cells were illuminated with Lambda DG-5 illumination system (Sutter Instruments, Novato, CA, USA), and imaging was performed using an acA720 camera (Basler, Ahrensburg, Germany) through a 40× oil immersion objective lens of an Olympus CKX53 inverted microscope. Taste bud cells loaded with Fura-2 AM were excited at 340 nm and 380 nm of light, and emission was recorded at 510 nm. Images were captured at a rate of 20 per minute, and the ratio of fluorescence (340 nm/380 nm) was used to measure the changes in intracellular calcium levels in taste bud cells by InCyt Im2™ imaging software (Version 6.00, Cincinnati, OH, USA).

### Nerve recording

Taste responsiveness in the chorda tympani (CT) nerve was recorded as previously described (Liu et al., 2021). Animals were anesthetized by intraperitoneal injection of urethane (about 2 g/kg), and tracheostomy was performed to facilitate breathing. The CT nerve was exposed through a ventral approach and incised near the tympanic bulla. A platinum-iridium electrode (A-M Systems) was attached to the severed nerve, and a reference electrode was placed in the nearby muscle for recording. A series of tastants, including sucrose (500 mM), NaCl (30 mM), capric acid (CA, 0.5 mM), denatonium benzoate (DB, 1 mM), HCl (1 mM), and intralipid (0.1, 0.5, 1, 2.5 and 5%), were applied for 15 second and rinsed with water for 45 seconds to the anterior tongue of the experimental animals. Model 3000 AC/DC differential amplifier (A-M Systems) and DataWave Technologies SciWorks Experimenter software (DataWave Technologies) were used for amplifying and collecting data. The area under the curve (AUC) of the nerve response for each tastant in WT and *Adipor1^-/-^* mice was calculated. NH_4_Cl (100 mM) was applied at the beginning and the end of each experiment, and taste responses were normalized to the average NH_4_Cl response within each session.

### Brief-access taste test

The brief-access testing was performed as described in previous study (Lin et al., 2024). Four groups of animals (*Adipor1^+/+^* males, *Adipor1^+/+^* females, *Adipor1^-/-^* males, and *Adipor1^-/-^* females; n = 8 for each group) were used. Before the test began, the mice were placed individually in standard cages and acclimated to the new environment for several days. The brief-access taste test was administered within an MS-160 Davis Rig gustatory behavioral apparatus (Med Associates, St. Albans, VT, USA). Water restriction was approximately 21 hours prior to each training and testing day (30-minute test, waiting for 1 hour, then free access to water for 1.5 hours). Purified water was used for testing and preparing a series of intralipid solutions (0.5, 1, 5, 10, and 20%). The well-trained animals were subjected to five days of testing of intralipid, a fan was placed near the Davis Rig chamber to provide constant airflow to reduce the olfactory attributes of the tastant. The test sessions were 30 minutes in duration, during which animals could initiate as many trials as possible. Due to the water deprivation, animals licked water or tastants profusely at the beginning, and once they became less thirsty, the number of water licks dropped dramatically compared to the preferred intralipid solutions. To better present the taste-associated differences in behavioral responses between water and intralipid, trials were included when the number of water licks began to drop dramatically, indicative of a reduction in thirst. The tastant/water lick ratio was calculated by dividing the mean number of licks per trial for each concentration by the mean number of water licks per trial for that animal.

### Conditioned taste aversion test

In the conditioned taste aversion (CTA) experiments, adult males and females from *Adipor1*^-/-^ and *Adipor1*^+/+^ mice were assigned into two categories. One is the experimental groups [*Adipor1*^-/-^ male (n = 8), *Adipor1*^-/-^ female (n = 6), *Adipor1*^+/+^ male (n = 7), and *Adipor1*^+/+^ female (n = 7)] in which taste aversions to 100 μM LA were conditioned with an intraperitoneal injection of 150 mM LiCl (20 ml/kg body weight), and the other is control groups [*Adipor1*^-/-^ male (n = 6), *Adipor1*^-/-^ female (n = 5), *Adipor1*^+/+^ male (n = 6), and *Adipor1*^+/+^ female (n = 8)] which was received 150 mM NaCl (20 ml/kg body weight). Details of the CTA assays have been described previously (Calder et al., 2021; Liu et al., 2021), including 3–4 days of water training, 3 days of conditioning, and 3 days of testing. Mice were deprived of water on the day before water training and started on a 22-hour water restriction schedule for the whole duration of the CTA assay. Animals were given 30 minutes of access to water each day, about 1 hour after the water training, conditioning, and testing to facilitate rehydration. An MS-160 Davis Rig gustatory behavioral apparatus (Med Associates) was used to perform water training and testing. A fan was placed near the Davis Rig chamber to provide constant airflow to reduce the olfactory component of the fatty acid. The solutions used in the CTA test are pure water, LA (1, 3, 10, 30, and 100 µM), sucrose (100 mM), and NaCl (75 mM). The sequences of the tastants were arranged randomly, and the test session included 2 blocks of 8 presentations with shutter opening for 5 seconds following 2 seconds of water rinse, and the wait times for the first lick were 150 s. The total number of licks per stimulus was averaged between the 2 trials and normalized relative to the water licks. Trials with zero licks, while infrequent, were removed from the analysis.

### Two-bottle preference assay

Animals caged individually were given several days of exposure to two sipper bottles containing water to habituate the mice to the experimental setting. In two-bottle preference assays, *Adipor1*^-/-^ mice and *Adipor1*^+/+^ controls were given access for 48 hours to two sipper bottles, one containing a test tastant solution and the other with water (n = 7–9 mice). The left-right positions of the tastant solution and water bottles were switched every 24 hours for each test to mitigate the effect of side preference. The sipper spouts and bottles were cleaned after every test and used randomly to avoid any association between tastant cues and spout cues. A 24-hour or 48-hour interval with two bottles of water was given between tastant solutions. The order of tastant solutions was as follows: LA (1, 3, 10, 30, and 100 µM), sucrose (30 mM), monosodium glutamate (MSG, 10 mM), HCl (1 mM), NaCl (75 mM), and DB (1 mM). Fluid intakes were measured by weighing the bottles at the beginning and the end of each 24-hour test. Intralipid (0.1%, 0.5%, 1%, 2.5%, and 5%) were applied for new groups of mice (n = 7–8 mice) and the intakes of intralipid were weighted and refilled every 12 hours. After the two-bottle preference test, these animals were used for other experiments, including the diet preference assay in Promethion system, high-fat diet feeding study, another two-bottle preference with intralipid, and nerve recording (timeline in Figure S5; *Supplement*). The ratio of 48-hour tastant intake over the total fluid (tastant + water) consumption was calculated as the preference ratio of the tastant.

### Promethion metabolic chambers

Promethion metabolic and behavioral cages (Sable Systems International, Las Vegas, NV) were utilized to compare the high-fat diet preference between *Adipor1*^-/-^ and WT mice. The Promethion system has a home-like environment with each cage containing a water bottle and two food hoppers, all connected to load cells for real-time weight monitoring. Animals (after the test of two-bottle preference for intralipid) were placed individually in cages with a chow diet in both food hoppers for four days (two days for adaptation to the new experimental environment, data collection in the following two days). The food in each hopper was then replaced with a control diet (10% calories from fat and 70% calories from carbohydrate) and a high-fat diet (60% calories from fat and 20% calories from carbohydrate). The position of the food hoppers was changed after 1 day of recording to avoid positional bias. The amount of 48-hour food consumption was measured and the preference ratio for the high-fat diet was calculated as high-fat diet intake over total food consumption. The system allows 16 animals to be tested at a time and we chose to test the males first and then the females (two female groups introduced in this experiment, n = 8 for each group).

### Feeding study

After removal from the Promethion cages, animals continued to be housed individually and had *ad libitum* access to high-fat diet and water. The animals’ body weights and the intakes of food and water were measured weekly. Total body composition (fat, lean tissue, and free body fluid) was determined by using the Bruker time domain-nuclear magnetic resonance (TD-NMR, minispec LF50) live mice body composition analyzer (Bruker, Billerica, MA, USA) before and after the high-fat diet study.

### Statistical analysis

Calcium imaging data analyses were based on the amplitude of the intracellular calcium concentration and analyzed in Origin 9.6 (Version 9.6.0.172, OriginLab, Northampton, MA, USA). Statistical analysis was performed using GraphPad Prism 10 (Version 10.2.1 (395), GraphPad Software, Boston, MA, USA). Unpaired student’s t-test, multiple unpaired t-tests (correct for multiple comparisons using the Holm-Šídák method), and two-way ANOVA were performed to compare the differences between the two genotypes. The main variable of interest was genotype effect. If a genotype effect was not observed, the interaction between genotype and concentration was explored. Multiple comparisons (Tukey’s) *post hoc* test was then conducted to determine which concentrations differed between the experimental groups. A web-based analysis tool (CalR) was used for the statistical analysis of energy expenditure and pedestrian locomotion data (Mina et al., 2018). The level of significance was set at α = 0.05 for all experiments. All data are presented as mean ± SEM.

## Acknowledgements

*Author Contributions:* F.L. and T.A.G. conceived and designed research; F.L. performed experiments; F.L. analyzed data; F.L. and T.A.G. interpreted results of experiments; F.L. and T.A.G. prepared figures; F.L. prepared first draft of manuscript; F.L. and T.A.G. edited and revised manuscript; F.L. and T.A.G. approved latest version of the manuscript; T.A.G. secured funding.

*Funding:* This research was supported by National Institutes of Health award R21DC021103 (tag).

*Institutional Review Board Statement:* All procedures involving animals were approved by the Institutional Animal Care and Use Committee of the University of Central Florida (protocol 2023-84; approved most recently on 4 June 2023) and were performed in accordance with American Veterinary Medical Association guidelines.

*Data* Availability *Statement:* All relevant data are included in the manuscript.

*Conflicts of Interest:* The authors declare no conflicts of interest, financial or otherwise.

## References

Ackroff, K., & Sclafani, A. (2014). Post-oral fat stimulation of intake and conditioned flavor preference in C57BL/6J mice: a concentration-response study. Physiology & Behavior, 129, 64–72.

Adamczak, M., Wiȩcek, A., Funahashi, T., Chudek, J., Kokot, F., & Matsuzawa, Y. (2003). Decreased plasma adiponectin concentration in patients with essential hypertension. American Journal of Hypertension, 16(1), 72–75.

Arita, Y., Kihara, S., Ouchi, N., Takahashi, M., Maeda, K., Miyagawa, J.-i., Hotta, K., Shimomura, I., Nakamura, T., & Miyaoka, K. (1999). Paradoxical decrease of an adipose-specific protein, adiponectin, in obesity. Biochemical and Biophysical Research Communications, 257(1), 79–83.

Bhattacharya, K., & Rattan, S. I. (2021). Fats and Oils for Health and Longevity. *Nutrition*, Food and Diet in Ageing and Longevity, 53–62.

Bjursell, M., Ahnmark, A., Bohlooly-Y, M., William-Olsson, L., Rhedin, M., Peng, X.-R., Ploj, K., Gerdin, A.-K., Arnerup, G., & Elmgren, A. (2007). Opposing effects of adiponectin receptors 1 and 2 on energy metabolism. Diabetes, 56(3), 583–593.

Bray, G. A., & Popkin, B. M. (1998). Dietary fat intake does affect obesity! The American Journal of Clinical Nutrition, 68(6), 1157–1173.

Brissard, L., Leemput, J., Hichami, A., Passilly-Degrace, P., Maquart, G., Demizieux, L., Degrace, P., & Khan, N. A. (2018). Orosensory detection of dietary fatty acids is altered in CB1R−/− mice. Nutrients, 10(10), 1347.

Cai, H., Cong, W.-n., Daimon, C. M., Wang, R., Tschöp, M. H., Sévigny, J., Martin, B., & Maudsley, S. (2013). Altered lipid and salt taste responsivity in ghrelin and GOAT null mice. PLoS One, 8(10), e76553.

Calder, A. N., Yu, T., Dahir, N. S., Sun, Y., & Gilbertson, T. A. (2021). Ghrelin Receptors Enhance Fat Taste Responsiveness in Female Mice. Nutrients, 13(4), 1045.

Calder, P. C. (2015). Functional roles of fatty acids and their effects on human health. Journal of Parenteral and Enteral Nutrition, 39, 18S–32S.

Chattopadhyay, S., Joharapurkar, A., Das, N., Khatoon, S., Kushwaha, S., Gurjar, A. A., Singh, A. K., Shree, S., Ahmed, M. Z., & China, S. P. (2022). Estradiol overcomes adiponectin-resistance in diabetic mice by regulating skeletal muscle adiponectin receptor 1 expression. Molecular and Cellular Endocrinology, 540, 111525.

Coope, A., Milanski, M., Araújo, E. P., Tambascia, M., Saad, M. J., Geloneze, B., & Velloso, L. A. (2008). AdipoR1 mediates the anorexigenic and insulin/leptin-like actions of adiponectin in the hypothalamus. FEBS Letters, 582(10), 1471–1476.

Crosson, S. M., Marques, A., Dib, P., Dotson, C. D., Munger, S. D., & Zolotukhin, S. (2019). Taste receptor cells in mice express receptors for the hormone adiponectin. Chemical Senses, 44(6), 409–422.

Dahir, N. S., Calder, A. N., McKinley, B. J., Liu, Y., & Gilbertson, T. A. (2021). Sex differences in fat taste responsiveness are modulated by estradiol. American Journal of Physiology-Endocrinology and Metabolism, 320(3), E566–E580.

Diószegi, J., Llanaj, E., & Ádány, R. (2019). Genetic background of taste perception, taste preferences, and its nutritional implications: a systematic review. Frontiers in Genetics, 10, 463497.

Dischinger, U., Corteville, C., Otto, C., Fassnacht, M., Seyfried, F., & Hankir, M. K. (2019). GLP-1 and PYY3-36 reduce high-fat food preference additively after Roux-en-Y gastric bypass in diet-induced obese rats. Surgery for obesity and related diseases, 15(9), 1483–1492.

Drewnowski, A. (1997). Taste preferences and food intake. Annual Review of Nutrition, 17(1), 237–253.

Drewnowski, A., & Greenwood, M. (1983). Cream and sugar: human preferences for high-fat foods. Physiology & Behavior, 30(4), 629–633.

Drewnowski, A., & Monsivais, P. (2020). Taste, cost, convenience, and food choices. In Present Knowledge in Nutrition (pp. 185-200). Elsevier.

Esmaili, S., Hemmati, M., & Karamian, M. (2020). Physiological role of adiponectin in different tissues: a review. Archives of Physiology and Biochemistry, 126(1), 67–73.

Gaillard, D., & Stratford, J. M. (2016). Measurement of behavioral taste responses in mice: Two-bottle preference, lickometer, and conditioned taste-aversion tests. Current Protocols in Mouse Biology, 6(4), 380–407.

Ghadge, A. A., Khaire, A. A., & Kuvalekar, A. A. (2018). Adiponectin: a potential therapeutic target for metabolic syndrome. Cytokine & Growth Factor Reviews, 39, 151–158.

Harnischfeger, F., & Dando, R. (2021). Obesity-induced taste dysfunction, and its implications for dietary intake. International Journal of Obesity, 45(8), 1644–1655.

Harris, R. B. (2019). Development of leptin resistance in sucrose drinking rats is associated with consuming carbohydrate-containing solutions and not calorie-free sweet solution. Appetite, 132, 114–121.

Karatayev, O., Baylan, J., & Leibowitz, S. F. (2009). Increased intake of ethanol and dietary fat in galanin overexpressing mice. Alcohol, 43(8), 571–580.

Kawai, K., Sugimoto, K., Nakashima, K., Miura, H., & Ninomiya, Y. (2000). Leptin as a modulator of sweet taste sensitivities in mice. Proceedings of the National Academy of Sciences, 97(20), 11044–11049.

Kitagawa, J., Shingai, T., Kajii, Y., Takahashi, Y., Taguchi, Y., & Matsumoto, S. (2007). Leptin modulates the response to oleic acid in the pharynx. Neuroscience Letters, 423(2), 109–112.

Kubota, N., Yano, W., Kubota, T., Yamauchi, T., Itoh, S., Kumagai, H., Kozono, H., Takamoto, I., Okamoto, S., & Shiuchi, T. (2007). Adiponectin stimulates AMP-activated protein kinase in the hypothalamus and increases food intake. Cell Metabolism, 6(1), 55–68.

Kumar, V., Shukla, A. K., Sharma, P., Choudhury, B., Singh, P., & Kumar, S. (2017). Role of macronutrient in health. World Journal of Pharmaceutical Research, 6(3), 373–381.

La Sala, M. S., Hurtado, M. D., Brown, A. R., Bohórquez, D. V., Liddle, R. A., Herzog, H., Zolotukhin, S., & Dotson, C. D. (2013). Modulation of taste responsiveness by the satiation hormone peptide YY. The FASEB Journal, 27(12), 5022–5033.

Lewandowski, D., Gao, F., Imanishi, S., Tworak, A., Bassetto, M., Dong, Z., Pinto, A. F., Tabaka, M., Kiser, P. D., & Imanishi, Y. (2024). Restoring retinal polyunsaturated fatty acid balance and retina function by targeting ceramide in AdipoR1 deficient mice. Journal of Biological Chemistry, 107291.

Li, X., Zhang, D., Vatner, D. F., Goedeke, L., Hirabara, S. M., Zhang, Y., Perry, R. J., & Shulman, G. I. (2020). Mechanisms by which adiponectin reverses high fat diet-induced insulin resistance in mice. Proceedings of the National Academy of Sciences, 117(51), 32584–32593.

Liem, D. G., & Russell, C. G. (2019). The influence of taste liking on the consumption of nutrient rich and nutrient poor foods. Frontiers in Nutrition, 6, 174.

Liman, E. R., & Kinnamon, S. C. (2021). Sour taste: receptors, cells and circuits. Current Opinion in Physiology, 20, 8–15.

Lin, F., Liu, Y., Rudeski-Rohr, T., Dahir, N., Calder, A., & Gilbertson, T. A. (2023). Adiponectin Enhances Fatty Acid Signaling in Human Taste Cells by Increasing Surface Expression of CD36. International Journal of Molecular Sciences, 24(6), 5801.

Liu, Y., Xu, H., Dahir, N., Calder, A., Lin, F., & Gilbertson, T. A. (2021). GPR84 Is Essential for the Taste of Medium Chain Saturated Fatty Acids. Journal of Neuroscience, 41(24), 5219–5228.

Loper, H. B., La Sala, M., Dotson, C., & Steinle, N. (2015). Taste perception, associated hormonal modulation, and nutrient intake. Nutrition Reviews, 73(2), 83–91.

Mack, C., Wilson, J., Athanacio, J., Reynolds, J., Laugero, K., Guss, S., Vu, C., Roth, J., & Parkes, D. (2007). Pharmacological actions of the peptide hormone amylin in the long-term regulation of food intake, food preference, and body weight. *American Journal of Physiology-Regulatory*, Integrative and Comparative Physiology, 293(5), R1855–R1863.

Maliphol, A. B., Garth, D. J., & Medler, K. F. (2013). Diet-induced obesity reduces the responsiveness of the peripheral taste receptor cells. PLoS One, 8(11), e79403.

Martin, B., Dotson, C. D., Shin, Y. K., Ji, S., Drucker, D. J., Maudsley, S., & Munger, S. D. (2009). Modulation of taste sensitivity by GLP-1 signaling in taste buds. Annals of the New York Academy of Sciences, 1170(1), 98–101.

Martin, C., Passilly-Degrace, P., Chevrot, M., Ancel, D., Sparks, S. M., Drucker, D. J., & Besnard, P. (2012). Lipid-mediated release of GLP-1 by mouse taste buds from circumvallate papillae: putative involvement of GPR120 and impact on taste sensitivity. Journal of Lipid Research, 53(11), 2256–2265.

Martin, C., Passilly-Degrace, P., Gaillard, D., Merlin, J.-F., Chevrot, M., & Besnard, P. (2011). The lipid-sensor candidates CD36 and GPR120 are differentially regulated by dietary lipids in mouse taste buds: impact on spontaneous fat preference. PLoS One, 6(8), e24014.

Masic, U., & Yeomans, M. R. (2017). Does acute or habitual protein deprivation influence liking for monosodium glutamate? Physiology & Behavior, 171, 79–86.

Mattes, R. D. (2021). Taste, teleology and macronutrient intake. Current Opinion in Physiology, 19, 162–167.

Meyerhof, W., Behrens, M., Brockhoff, A., Bufe, B., & Kuhn, C. (2005). Human bitter taste perception. Chemical Senses, 30(suppl_1), i14-i15.

Mina, A. I., LeClair, R. A., LeClair, K. B., Cohen, D. E., Lantier, L., & Banks, A. S. (2018). CalR: a web-based analysis tool for indirect calorimetry experiments. Cell Metabolism, 28(4), 656–666. e651.

Mizushige, T., Inoue, K., & Fushiki, T. (2007). Why is fat so tasty? Chemical reception of fatty acid on the tongue. Journal of Nutritional Science and Vitaminology, 53(1), 1–4.

Mullen, K. L., Pritchard, J., Ritchie, I., Snook, L. A., Chabowski, A., Bonen, A., Wright, D., & Dyck, D. J. (2009). Adiponectin resistance precedes the accumulation of skeletal muscle lipids and insulin resistance in high-fat-fed rats. *American Journal of Physiology-Regulatory*, Integrative and Comparative Physiology, 296(2), R243–R251.

Mullen, K. L., Smith, A. C., Junkin, K. A., & Dyck, D. J. (2007). Globular adiponectin resistance develops independently of impaired insulin-stimulated glucose transport in soleus muscle from high-fat-fed rats. American Journal of Physiology-Endocrinology and Metabolism, 293(1), E83–E90.

Niki, M., Jyotaki, M., Yoshida, R., Yasumatsu, K., Shigemura, N., DiPatrizio, N. V., Piomelli, D., & Ninomiya, Y. (2015). Modulation of sweet taste sensitivities by endogenous leptin and endocannabinoids in mice. The Journal of Physiology, 593(11), 2527–2545.

Okada, S., York, D., Bray, G., & Erlanson-Albertsson, C. (1991). Enterostatin (Val-Pro-Asp-Pro-Arg), the activation peptide of procolipase, selectively reduces fat intake. Physiology & Behavior, 49(6), 1185–1189.

Pfabigan, D. M., Vezzani, C., Thorsby, P. M., & Sailer, U. (2022). Sex difference in human olfactory sensitivity is associated with plasma adiponectin. Hormones and Behavior, 145, 105235.

Pittman, D. W., Smith, K. R., Crawley, M. E., Corbin, C. H., Hansen, D. R., Watson, K. J., & Gilbertson, T. A. (2008). Orosensory detection of fatty acids by obesity-prone and obesity-resistant rats: strain and sex differences. Chemical Senses, 33(5), 449–460.

Primeaux, S. D., York, D. A., & Bray, G. A. (2006). Neuropeptide Y administration into the amygdala alters high fat food intake. Peptides, 27(7), 1644–1651.

Qi, Y., Takahashi, N., Hileman, S. M., Patel, H. R., Berg, A. H., Pajvani, U. B., Scherer, P. E., & Ahima, R. S. (2004). Adiponectin acts in the brain to decrease body weight. Nature Medicine, 10(5), 524–529.

Salmeron, J., Hu, F. B., Manson, J. E., Stampfer, M. J., Colditz, G. A., Rimm, E. B., & Willett, W. C. (2001). Dietary fat intake and risk of type 2 diabetes in women. The American Journal of Clinical Nutrition, 73(6), 1019–1026.

Shigemura, N., Ohta, R., Kusakabe, Y., Miura, H., Hino, A., Koyano, K., Nakashima, K., & Ninomiya, Y. (2004). Leptin modulates behavioral responses to sweet substances by influencing peripheral taste structures. Endocrinology, 145(2), 839–847.

Shimbara, T., Mondal, M. S., Kawagoe, T., Toshinai, K., Koda, S., Yamaguchi, H., Date, Y., & Nakazato, M. (2004). Central administration of ghrelin preferentially enhances fat ingestion. Neuroscience Letters, 369(1), 75–79.

Shin, Y. K., Martin, B., Golden, E., Dotson, C. D., Maudsley, S., Kim, W., Jang, H. J., Mattson, M. P., Drucker, D. J., & Egan, J. M. (2008). Modulation of taste sensitivity by GLP-1 signaling. Journal of Neurochemistry, 106(1), 455–463.

Sieri, S., Krogh, V., Ferrari, P., Berrino, F., Pala, V., Thiébaut, A. C., Tjønneland, A., Olsen, A., Overvad, K., & Jakobsen, M. U. (2008). Dietary fat and breast cancer risk in the European Prospective Investigation into Cancer and Nutrition. The American Journal of Clinical Nutrition, 88(5), 1304–1312.

Smith, B. K., Andrews, P. K., & West, D. B. (2000). Macronutrient diet selection in thirteen mouse strains. *American Journal of Physiology-Regulatory*, Integrative and Comparative Physiology, 278(4), R797–R805.

Stewart, J., & Keast, R. (2012). Recent fat intake modulates fat taste sensitivity in lean and overweight subjects. International Journal of Obesity, 36(6), 834–842.

Swinburn, B. A., Sacks, G., Hall, K. D., McPherson, K., Finegood, D. T., Moodie, M. L., & Gortmaker, S. L. (2011). The global obesity pandemic: shaped by global drivers and local environments. The Lancet, 378(9793), 804–814.

Treesukosol, Y., & Moran, T. H. (2022). Administration of Exendin-4 but not CCK alters lick responses and trial initiation to sucrose and intralipid during brief-access tests. Chemical Senses, 47, bjac004.

Tung, Y. L., Rimmington, D., O’Rahilly, S., & Coll, A. P. (2007). Pro-opiomelanocortin modulates the thermogenic and physical activity responses to high-fat feeding and markedly influences dietary fat preference. Endocrinology, 148(11), 5331–5338.

Ullah, H., Khan, A. S., Murtaza, B., Hichami, A., & Khan, N. A. (2021). Tongue leptin decreases oro-sensory perception of dietary fatty acids. Nutrients, 14(1), 197.

Wang, J.-J., Liang, K.-L., Lin, W.-J., Chen, C.-Y., & Jiang, R.-S. (2020). Influence of age and sex on taste function of healthy subjects. PLoS One, 15(6), e0227014.

Yamauchi, T., Kamon, J., Waki, H., Terauchi, Y., Kubota, N., Hara, K., Mori, Y., Ide, T., Murakami, K., & Tsuboyama-Kasaoka, N. (2001). The fat-derived hormone adiponectin reverses insulin resistance associated with both lipoatrophy and obesity. Nature Medicine, 7(8), 941–946.

Yamauchi, T., Nio, Y., Maki, T., Kobayashi, M., Takazawa, T., Iwabu, M., Okada-Iwabu, M., Kawamoto, S., Kubota, N., & Kubota, T. (2007). Targeted disruption of AdipoR1 and AdipoR2 causes abrogation of adiponectin binding and metabolic actions. Nature Medicine, 13(3), 332–339.

Yoshida, R., Margolskee, R. F., & Ninomiya, Y. (2021). Phosphatidylinositol-3 kinase mediates the sweet suppressive effect of leptin in mouse taste cells. Journal of Neurochemistry, 158(2), 233–245.

Yoshida, R., Noguchi, K., Shigemura, N., Jyotaki, M., Takahashi, I., Margolskee, R. F., & Ninomiya, Y. (2015). Leptin suppresses mouse taste cell responses to sweet compounds. Diabetes, 64(11), 3751–3762.

Yoshida, R., Ohkuri, T., Jyotaki, M., Yasuo, T., Horio, N., Yasumatsu, K., Sanematsu, K., Shigemura, N., Yamamoto, T., & Margolskee, R. F. (2010). Endocannabinoids selectively enhance sweet taste. Proceedings of the National Academy of Sciences, 107(2), 935–939.

Zeng, W., Jin, Q., & Wang, X. (2023). Reassessing the Effects of Dietary Fat on Cardiovascular Disease in China: A Review of the Last Three Decades. Nutrients, 15(19), 4214.

Zhao, J., Lyu, C., Gao, J., Du, L., Shan, B., Zhang, H., Wang, H.-Y., & Gao, Y. (2016). Dietary fat intake and endometrial cancer risk: A dose response meta-analysis. Medicine, 95(27), e4121.

