## Supplementary Material for "Fat taste responsiveness, but not dietary fat intake, is affected in *Adipor1* null mice"

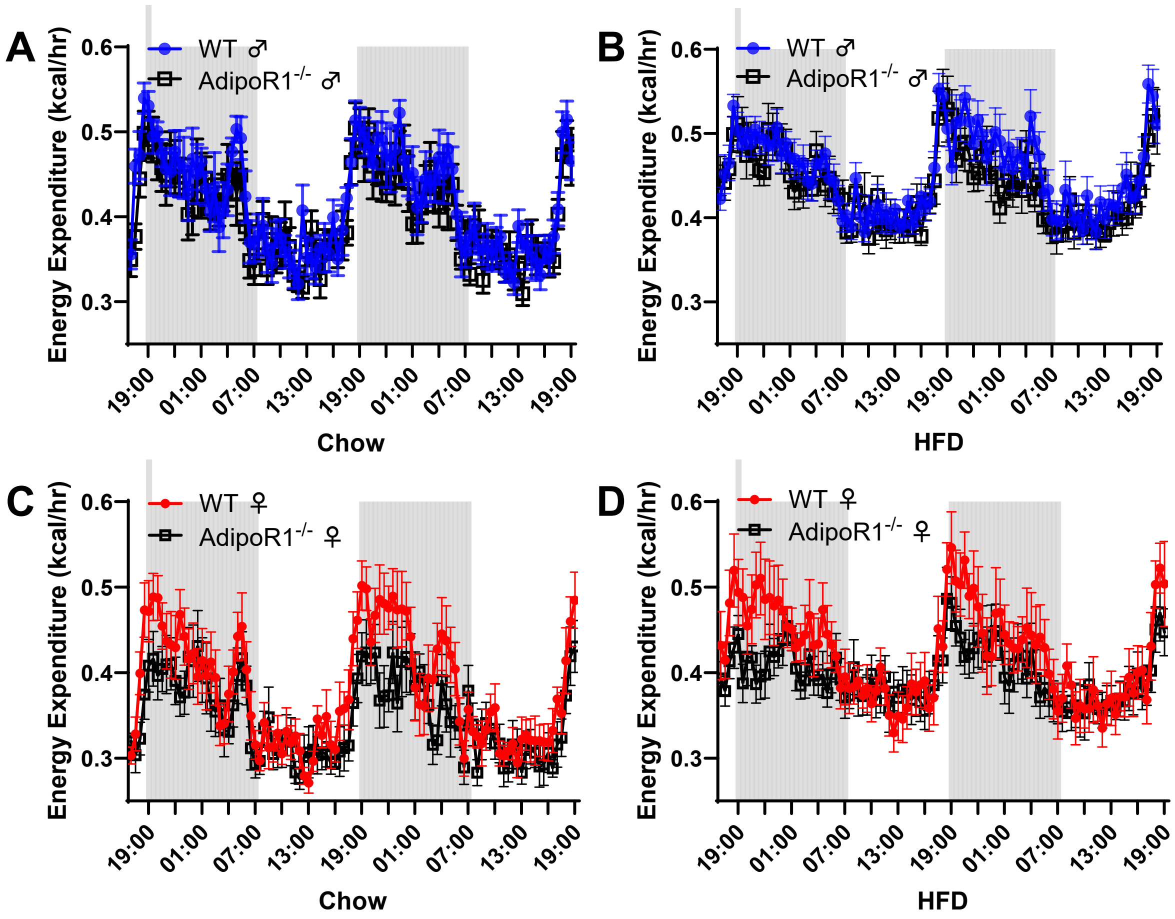


**Figure S1.** Energy expenditure in *Adipor1*$^{-/-}$ and WT mice on chow and high-fat diets. No significant difference in energy expenditure was found between *Adipor1*$^{-/-}$ and WT males on a chow (A) or a high-fat diet (B); Reduced energy expenditure were seen in *Adipor1*$^{-/-}$ females at some time points during the dark phase (gray shaded), but no overall significant differences were found between *Adipor1*$^{-/-}$ and WT females on chow (C) and high-fat diets (D). A web-based analysis tool (CalR) was used for the statistical analysis. Data are presented as mean ± SEM (n = 8 per group).

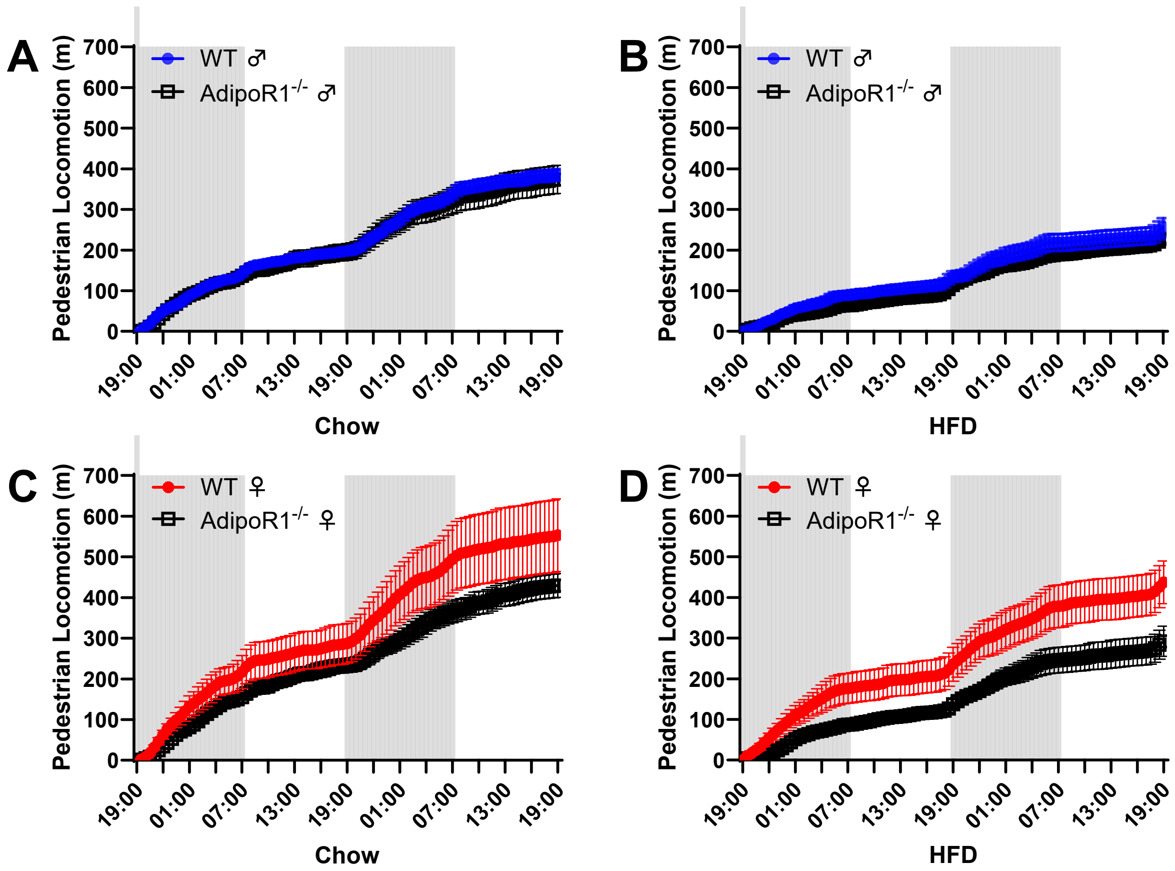


**Figure S2.** Pedestrian locomotion in *Adipor1*$^{-/-}$ and WT mice on chow and high-fat diets. (A) No significant difference in pedestrian locomotion was found between *Adipor1*$^{-/-}$ and WT males on a chow diet; (B) *Adipor1*$^{-/-}$ males showed a slightly greater decrease in pedestrian locomotion compared to WT males on a high-fat diet; (C) *Adipor1*$^{-/-}$ females showed a slight decrease in pedestrian locomotion compared to WT females on a chow diet; (D) *Adipor1*$^{-/-}$ females showed a significant decrease in pedestrian locomotion compared to WT females on a high-fat diet ($P<0.05$). A web-based analysis tool (CalR) was used for the statistical analysis. Data are presented as mean ± SEM (n = 8 per group).

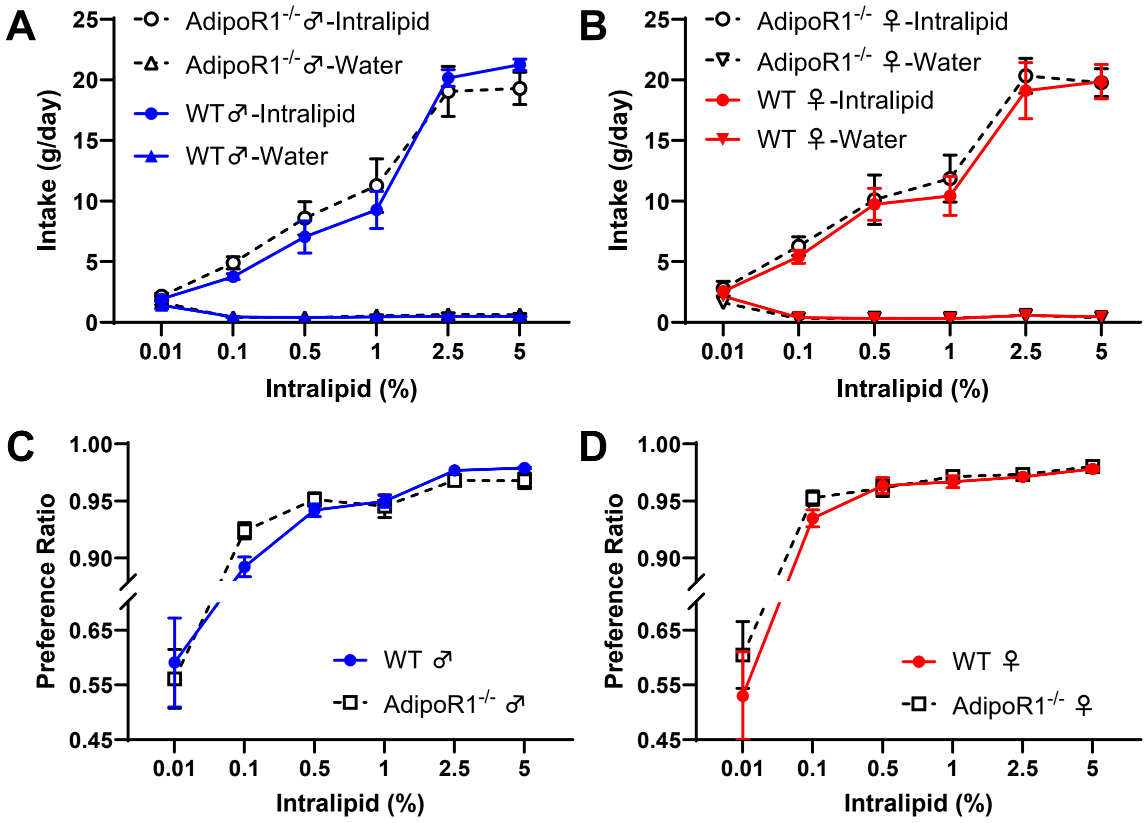


**Figure S3.** The intake and preference of different concentrations of intralipid in high-fat diet-fed *Adipor1*$^{-/-}$ and WT mice. No significant genotype effect and interaction were observed in intralipid consumption in both males (A) and females (B); There was no significant difference in intralipid preference between *Adipor1*$^{-/-}$ and WT mice in both males (C) and females (D). The ratio of 48-hour tastant intake over the total fluid (tastant + water) consumption was calculated as the preference ratio of the tastant. Data are presented as mean ± SEM (n = 7–8). Two-way ANOVA was performed to compare the differences between the two genotypes, with Tukey's multiple comparisons *post hoc* test conducted to determine which intralipid concentrations differed.

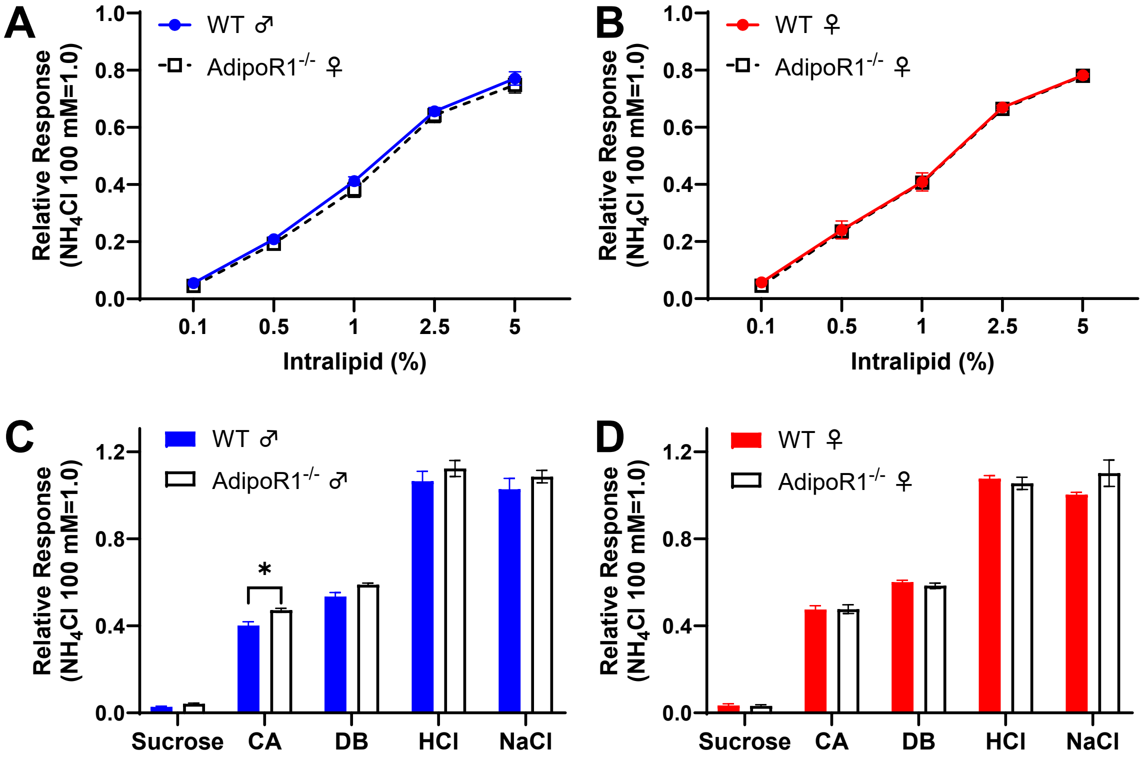


**Figure S4.** Chorda tympani (CT) nerve responses to different concentrations of intralipid and various tastants (15 s) in high-fat diet-fed *Adipor1*$^{-/-}$ and WT mice. Dose-dependent CT nerve responses to intralipid were observed in all animal groups, and there was no significant genotype effect on these nerve responses in males (A) and females (B); (C) Male *Adipor1*$^{-/-}$ mice displayed a significantly increased nerve response to capric acid (CA, 0.5 mM, Adjusted P=0.025) and no difference in nerve responses to sucrose (500 mM), denatonium benzoate (DB, 1 mM), HCl (1 mM), and NaCl (30 mM) was observed. (D) No significant differences were observed between the two genotype animals in females when stimulated with sucrose (500 mM), CA (0.5 mM), DB (1 mM), HCl (1 mM), and NaCl (30 mM). $NH_4Cl$ (100 mM) was applied at the beginning and the end of each experiment, and taste responses were normalized to the average $NH_4Cl$ response within each session. The area under the curve (AUC) of the nerve response for each tastant was calculated. Responses show the mean ± SEM (n = 4–6 mice). Two-way ANOVA was performed to compare the differences between the two genotypes, with Tukey's multiple comparisons *post hoc* test was conducted to determine which intralipid concentrations differed. Multiple unpaired t-tests, correct for multiple comparisons using the Holm-Šídák method, were used to compare the other tastants between the two genotypes. * $p < 0.05$.

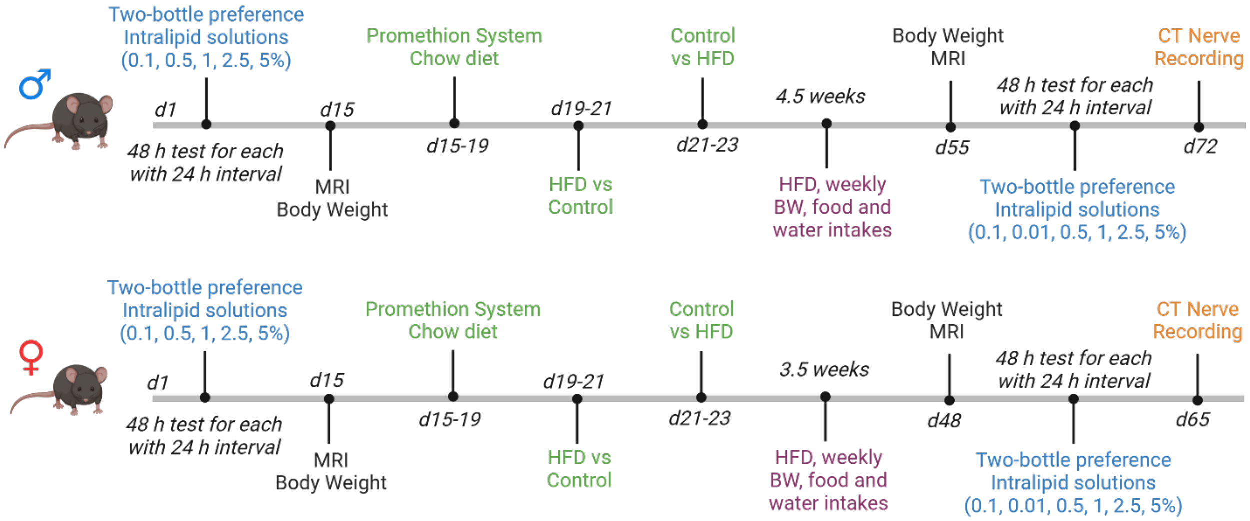


**Figure S5.** Timeline and design of the experiment. *Adipor1*$^{-/-}$ male (n = 8), *Adipor1*$^{+/+}$ male (n = 8), *Adipor1*$^{-/-}$ female (n = 7), and *Adipor1*$^{+/+}$ female (n = 7) entered the study at approximately 9 weeks of age. After 1 week of acclimation to the experimental setting, all the animals were performed with a two-bottle preference test for intralipid (as the order of 0.1, 0.5, 1, 2.5, and 5%) starting on day 1 (d1) in the next 2 weeks. Before the diet preference in the Promethion system, the body weight and body composition were measured. The Promethion system only allows 16 animals to be tested at a time and we chose to test the males first and then the females. Animals were placed individually in cages with chow diet in both food hoppers for four days (adaptation to the new experimental environment in the first two days, data analysis in the next two days). The food in each food hopper was then replaced with a control diet and a high-fat diet. The position of the food hoppers was changed after 1 day of recording to avoid position bias. Next, animals were given a high-fat diet feeding study, and body weight, food and water intakes were measured weekly. The body weight and body composition were measured again at the end of feeding study. High-fat diet fed animals received chow diet for several days, and we then subjected these animals to another two-bottle preference test for intralipid (as the order of 0.1, 0.01, 0.5, 1, 2.5, and 5%), followed by CT nerve recordings. Created in https://BioRender.com
